# The mitochondrial ferredoxin-like is essential for the formation of complex I-containing respiratory supercomplexes in *Arabidopsis thaliana*

**DOI:** 10.1101/2022.09.02.506396

**Authors:** Helene Röhricht, Jonathan Przybyla-Toscano, Joachim Forner, Clément Boussardon, Olivier Keech, Nicolas Rouhier, Etienne H. Meyer

## Abstract

In eukaryotes, mitochondrial ATP is mainly produced by the oxidative phosphorylation (OXPHOS) system, which is composed of five multiprotein complexes (complexes I to V). Analyses of the OXPHOS system by native gel electrophoresis revealed an organization of OXPHOS complexes in supercomplexes, but their roles and assembly pathways remain unclear. In this study, we characterized an atypical mitochondrial ferredoxin (mFDX-like). This protein was previously found associated with complex I, being part of the bridge domain linking the matrix and membrane arms of the complex. A phylogenetic analysis suggests that the *Arabidopsis thaliana* mFDX-like evolved from classical mitochondrial ferredoxin, but it lost one of the cysteines required for the coordination of the iron-sulfur (Fe-S) cluster essential for the electron transfer function of ferredoxins. Accordingly, our biochemical study shows that AtmFDX-like does not bind an Fe-S cluster, and is therefore unlikely to be involved in electron transfer reactions. To study the function of mFDX-like, we created deletion lines in Arabidopsis using a CRISPR/Cas9 approach. These lines do not show any growth phenotype under standard growth condition. However, the characterization of the OXPHOS system demonstrates that mFDX-like is important for the assembly of complex I, and essential for the formation of complex I-containing supercomplexes. We propose that mFDX-like and the bridge domain are required for the correct conformation of the membrane arm of complex I that is essential for the association of complex I with complex III to form supercomplexes.

## Introduction

Respiration is a metabolic pathway essential for plant growth as it produces energy, in the form of ATP, required to sustain metabolism (Nicholls and Ferguson, 2013). Respiratory ATP is mainly produced in mitochondria by the ATP synthase located in the inner mitochondrial membrane (IMM). To convert ADP into ATP, the ATP synthase utilizes the electrochemical gradient built across the IMM by the respiratory chain. The respiratory chain recycles cofactors for upstream metabolic reactions and transfers protons from the matrix to the intermembrane space. Together the respiratory chain and the ATP synthase form the oxidative phosphorylation (OXPHOS) system. The plant OXPHOS system comprises five protein complexes (named complex I to V, complex V being the ATP synthase) and several individual proteins as well as a mobile electron carrier coenzyme Q (Braun, 2020).

The biochemical analysis of OXPHOS complexes usually involves native gel electrophoresis. After isolating mitochondria, protein complexes are solubilized using mild detergents and separated on a native gel (Schagger and von Jagow, 1991). Using very mild detergents such as digitonin or triton X100, additional bands, larger than the individual complexes, are observed and further analyses determined that these bands correspond to assemblies of OXPHOS complexes (Schagger and Pfeiffer, 2001). These stoichiometric arrangements of OXPHOS complexes are called supercomplexes. Such assemblies correspond to homodimers (complex V dimers), heterodimers (supercomplexes I+III or III+IV) or larger combinations (e.g. supercomplex I+III+IV) (Lenaz and Genova, 2009, Enriquez, 2016). Supercomplexes were observed in all organisms investigated so far. However, the roles of these arrangements remain unresolved (Milenkovic et al., 2017, Hirst, 2018, Wu et al., 2020, Stuchebrukhov et al., 2020, Javadov et al., 2021).

Complex I is the largest OXPHOS complex (Hirst, 2013). In plants, it is composed of at least 47 subunits, of which nine subunits are encoded by genes present in the mitochondrial genome (Braun et al., 2014, Peters et al., 2013). Complex I is composed of two arms, the membrane arm embedded in the IMM, and the matrix arm that is attached at one end of the membrane arm. Each arm is composed of two modules. The matrix arm is formed by the N module where NADH binds and the Q module where ubiquinone is reduced. The membrane arm is divided into the so-called proximal (P_P_) module, where the matrix arm is bound and the distal (P_D_) module forming the tip of the membrane arm (Letts and Sazanov, 2015, Efremov and Sazanov, 2012). In plants, additional structures are present on the matrix side of the membrane arm: a globular structure formed by gamma carbonic anhydrases (CA), the CA domain and a bridge domain linking the matrix arm and the CA domain (Klusch et al., 2021, Soufari et al., 2020, Maldonado et al., 2020). Genes encoding subunits of the CA domain are present in the genomes of many eukaryotes but not opisthokonts, suggesting that this feature was present in LECA but has been lost in fungi and animals (Cainzos et al., 2021). Complex I oxidizes NADH at the tip of the matrix arm and transfers electrons via several iron-sulfur (Fe-S) clusters across this arm to reduce ubiquinone at the interface between the matrix and membrane arms (Lopez-Lopez et al., 2022). Concomitantly, protons are transferred from the matrix to the IMS by the membrane arm via an uncharacterized mechanism (Baradaran et al., 2013, Kampjut and Sazanov, 2020, Kolata and Efremov, 2021, Parey et al., 2021). In plants, complex I activity is essential as null mutants are unable to establish a seedling (Kühn et al., 2015) or are male-sterile (Gutierres et al., 1997).

Complex I is one of the most studied OXPHOS complexes in plants (Braun et al., 2014); yet many aspects of its biology remain unclear. For example, its composition is not fully resolved as structural work indicates that at least one additional subunit would be present (Klusch et al., 2021, Soufari et al., 2020, Maldonado et al., 2020). In addition, little is known about the regulation of complex I activity *in planta*. In order to identify genes encoding mitochondrial proteins playing important roles for complex I function, we performed a gene function prediction approach using EnsembleNet (Hansen et al., 2018). One of the identified candidate genes encodes a mitochondrial protein belonging to the ferredoxin (FDX) family.

In eukaryotes, FDXs are conserved cysteinyl-ligated [2Fe-2S] proteins participating to a wide range of oxidation-reduction reactions. In plants, FDXs are present both in plastids and mitochondria (Takubo et al., 2003, Przybyla-Toscano et al., 2018). The chloroplastic isoforms are at a metabolic crossroad, controlling the electron flow towards CO_2_ fixation, nitrogen and sulfur assimilation or lipid desaturation for instance (Przybyla-Toscano et al., 2018, Hanke and Mulo, 2013). Concerning mitochondrial representatives, mFDX1 and mFDX2 have been associated with cofactor (*i*.*e*. biotin) and hormone (*i*.*e*. homocastasterone) biosynthetic pathways in Arabidopsis (Picciocchi et al., 2003, Bellido et al., 2022); and more importantly they should be involved in the *de novo* synthesis of Fe-S clusters by analogy with the model in other organisms (Przybyla-Toscano et al., 2021). They receive electrons from a mitochondrial NADPH-dependent FDX reductase (FDXR, also known as adrenodoxin reductase in animals). Functional analysis of *fdxr* mutants and double mutants *mfdx1 mfdx2* in *Arabidopsis thaliana* revealed that the function of this FDXR-FDX system is critical for female gametophyte development and for early embryogenesis through a maternal effect (Bellido et al., 2022). In *A. thaliana*, a third gene encoding a putative FDX (At3g07480) has been identified in mitochondrial proteome studies (Heazlewood et al., 2004, Senkler et al., 2017, Niehaus et al., 2020, Fuchs et al., 2020). Interestingly, in a recently released cryo-EM complex I structure from Arabidopsis, this protein was identified as a structural subunit present in the bridge domain linking the matrix arm and the CA domain, together with an acyl carrier protein (ACP) (Klusch et al., 2021). However, one cysteine that normally serves as Fe-S cluster ligand in classical FDXs is absent, and accordingly only a metal ion was observed in the structure despite it adopts the overall fold of FDX. Moreover, functional studies have not been performed for this protein *in planta*.

In this study, we performed an *in vitro* biochemical characterization of the recombinant Arabidopsis mFDX-like (AtmFDX-like), which demonstrated that mFDX-like does not coordinate an Fe-S cluster unless the missing cysteine is reincorporated. This suggests that mFDX-like is not involved in electron transfer reactions. Using CRISPR-Cas9 lines, we also showed that an Arabidopsis *mfdx-like* knockout mutant does not exhibit any growth phenotype when grown under standard conditions. However, at the molecular level, we evidenced that mFDX-like is essential for the formation of complex I-containing respiratory supercomplexes in plant mitochondria.

## Results

### The atypical mFDX-like is present in several eukaryotic groups of the green lineage

To delineate the function of the AtmFDX-like, we firstly analyzed and compared its protein sequence with the ones of mitochondrial FDX orthologs from Arabidopsis, yeast and human (Supplemental Figure S1). Alignment of primary sequences revealed that Arabidopsis mFDX-like represents an atypical isoform sharing only approximately 20% sequence identity with Arabidopsis mFDX1/2, whereas Arabidopsis mtFDX1 and 2 share 76% identity even when including their targeting sequences. Assuming that all these sequences diverged from a single ancestor gene, mFDX-like diverged significantly. In particular and as already mentioned, one of the four cysteines serving as ligands for the putative Fe-S cluster is absent, and two other cysteines are separated by three amino acids instead of two in regular FDXs. Moreover, most negatively charged residues known to be involved in the interaction with the FDXR are not conserved, raising questions about its function(s) and partner(s). Standard PSI-BLAST searches using AtmFDX proteins as queries identified both classical and atypical mFDXs, despite the fact that mFDX-like proteins form a clade diverging from classical mFDXs, and obviously from plastidial FDXs (Figure 1). A careful inspection of mFDX-like distribution revealed that mFDX-like proteins are restricted to eukaryotes of the green lineage as demonstrated by their absence in cyanobacteria but presence in some algae, notably chlorophytes, as well as in bryophytes, lycophytes and angiosperms. The NUOP3 protein from *Chlamydomonas reinhardtii* that was previously shown to be also associated with complex I (Cardol, 2011) is in the same branch as AtmFDX-like (Figure 1) despite that they have diverged substantially, since both proteins have 20% identity (Supplemental Figure S2). Worth mentioning, most algal orthologs do not possess the three remaining cysteines known to serve as Fe-S cluster ligands in regular FDXs, whereas they are still present in orthologs from terrestrial plants. This suggests their progressive loss during evolution and dispensability at least for a role as a complex I structural subunit. Intriguingly, the NDUFX protein present in the complex I structure from the free-living protozoa *Tetrahymena thermophila* belonging to the Ciliophora phylum (Zhou et al., 2022) is present in the same phylogenetic clade (Figure 1).

**Figure 1.**
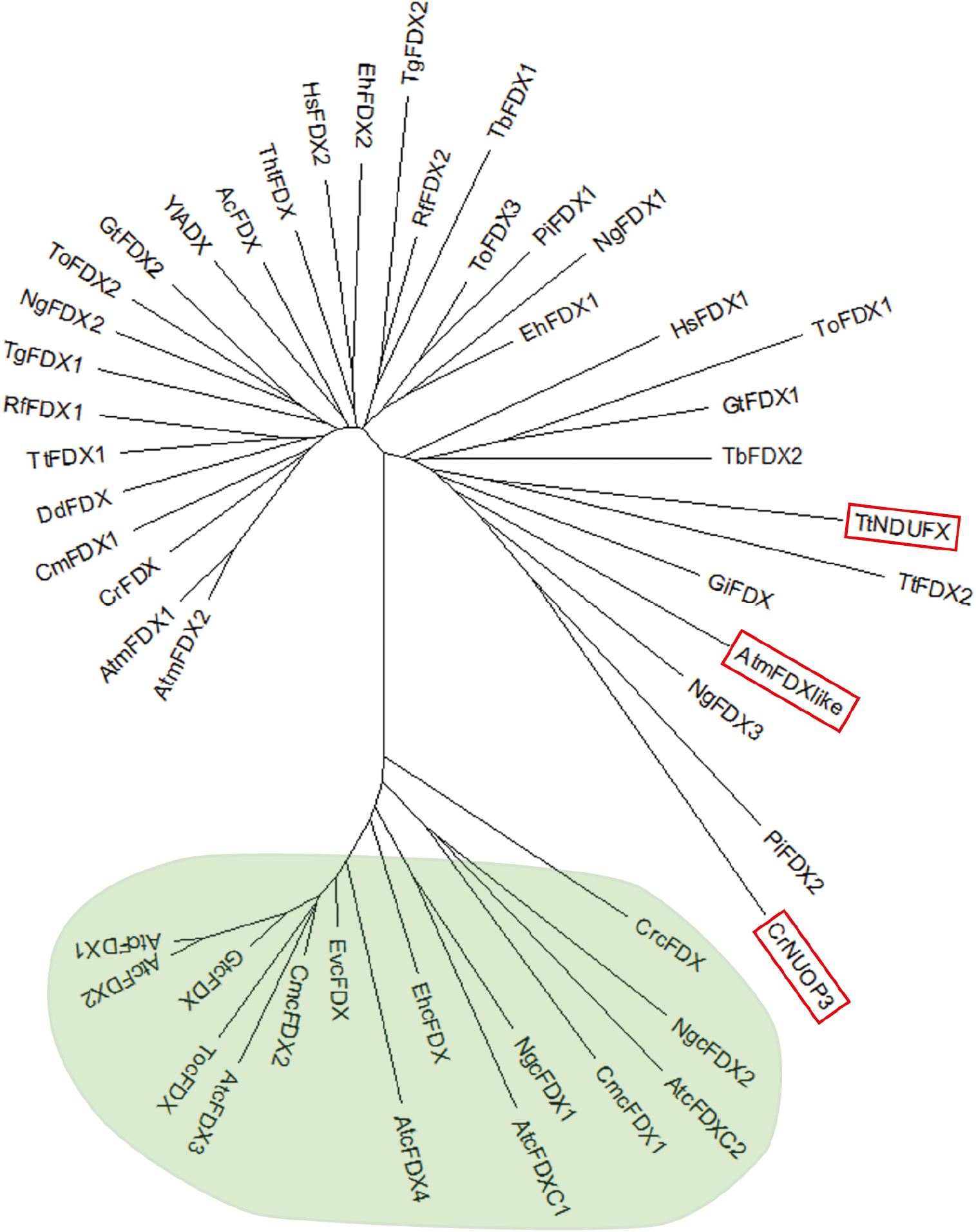
The atypical mFDX-like proteins form a phylogenetic branch distinct from other eukaryotic FDXs. Protein sequences of FDXs were retrieved from a selection of eukaryotic organisms representing the different eukaryotic lineages (see Supplemental Table S1). Sequences were analyzed using MEGA (www.megasoftware.net) and a minimum evolution tree was built using default parameters. Plastid FDXs are circled in green. The three FDXs described as complex I subunits are shown with red boxes. Ac: *Acanthamoeba castellanii*, At: *Arabidopsis thaliana*, Cm: *Cyanidioschyzon merolae*, Cr: *Chlamydomonas reinhardtii*, Dd: *Dictyostelium discoideum*, Eh: *Emiliania huxleyi*, Ev: *Euglena viridis*, Gi: *Giardia intestinalis*, Gt: *Guillardia theta*, Hs: *Homo sapiens*, Ng: *Nannochloropsis gaditana*, Pi: *Phytophtora infestans*, Rf: *Reticulomyxa filosa*, Tb: *Trypanosoma brucei*, Tg: *Toxoplasma gondii*, Th: *Thecamonas trahens*, To: *Thalassiosira oceanica*, Tt: *Tetrahymena thermophila*, Yl: *Yarrowia lipolytica*. The UniProt accessions of all the proteins analyzed are given in Supplemental Table S1.

### The absence of a single cysteine residue prevents mFDX-like to bind the regular [2Fe-2S] cluster

From the 3D structure analysis of Arabidopsis mFDX-like, it was noticed that a histidine residue, which may serve as an Fe-S cluster ligand in some proteins, is positioned in the region where the cysteine ligand normally is. In order to obtain clear evidence that this protein indeed lost the capacity to bind the [2Fe-2S] cluster normally present in classical mFDXs, we individually expressed the three Arabidopsis mFDXs in an *Escherichia coli* heterologous system, and purified the corresponding recombinant proteins (Supplemental Figure S3). After purification, only mFDX1 and mFDX2 exhibited a strong reddish-brown coloration. Accordingly, their UV-visible absorption spectra showed characteristic absorption bands at 342, 416, and 457 nm typical of the presence of a [2Fe-2S] cluster (Figure 2A and 2B). In contrast, no similar absorption bands were visible for mFDX-like (Figure 2C), indicating that the histidine residue cannot substitute for the cysteine ligand. To understand whether more pronounced structural changes occurred or whether introducing a single cysteine residue is enough for restoring the [2Fe-2S] cluster binding capacity, we have also expressed a variant of mFDX-like in which the leucine at position 85 was replaced by a cysteine (L85C). The purified mutated variant showed an absorption spectrum comprising shoulders centered at 333, 414 and 452 nm, very similar to that observed for the [2Fe-2S] cluster-bound forms of mFDX1 and mFDX2 (Figure 2D). Altogether, these *in vitro* analyses indicate that the loss of a cysteine ligand hampers [2Fe-2S] cluster binding by mFDX-like but that the protein still possesses the structural capacity to accommodate it when the cysteine is re-incorporated. In line with this, no [2Fe-2S] cluster was observed in Arabidopsis mFDX-like present in complex I structure but instead a single metal ion was coordinated by the residual cysteines and the histidine residue (Klusch et al., 2021).

**Figure 2.**
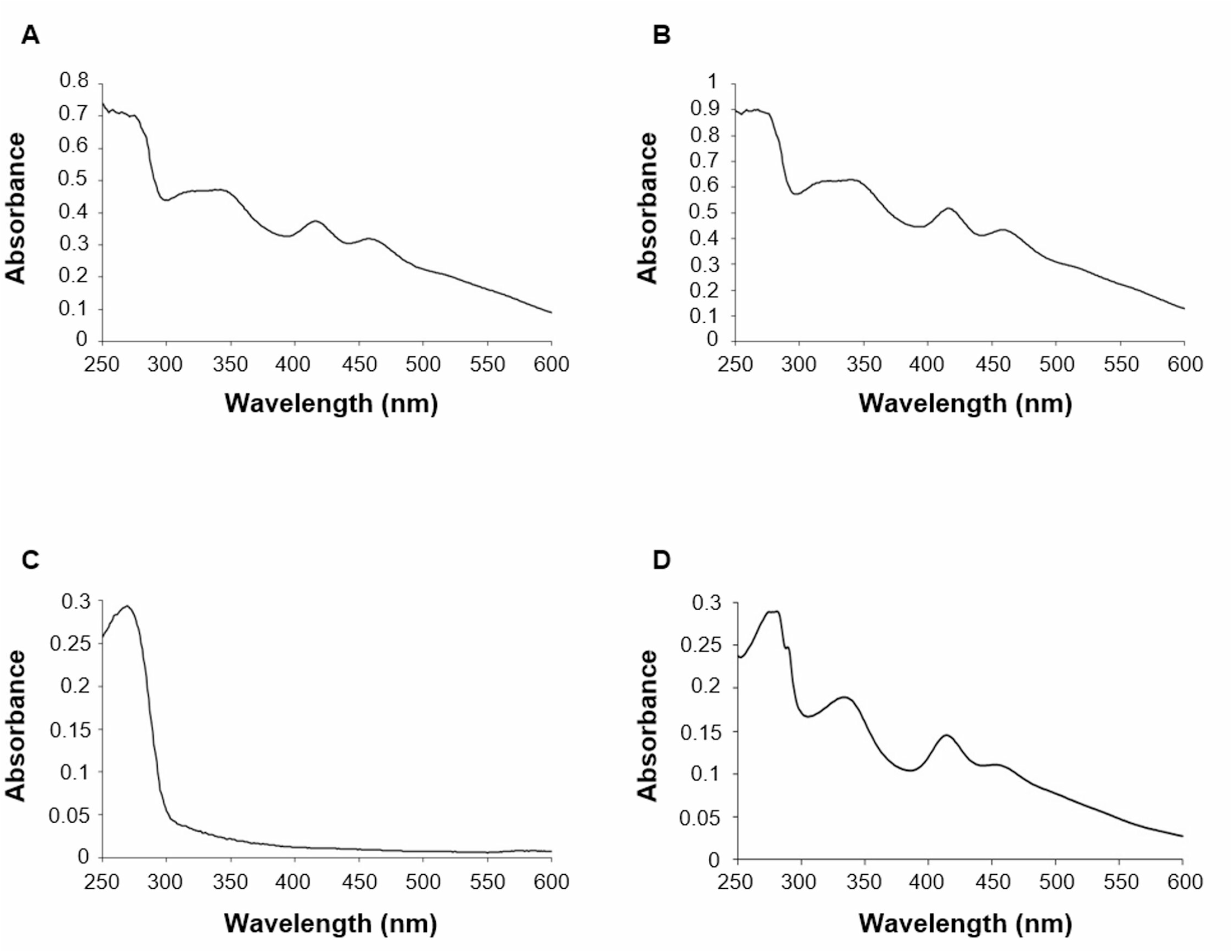
mFDX-like does not coordinate an Fe-S cluster. UV-visible absorption spectra of as-purified *Arabidopsis thaliana* his-tagged mFDX1 (A), mFDX2 (B), mFDX-like (C) and mFDX-like L85C variant (D) recombinantly expressed in *E. coli*.

### mFDX-like is a mitochondrial protein

mFDX-like is present in mitochondrial proteomes and in complex I structure (Klusch et al., 2021). To confirm that it is localized only in mitochondria *in vivo*, we generated stable Arabidopsis lines expressing a mFDX-GFP construct, and assessed its subcellular distribution by confocal laser microscopy. We observed that the mFDX-like-GFP signal colocalized nearly exclusively with the MitoTraker in root cells from 14-day-old seedlings, strengthening a “mitochondrion-only” targeting of the protein (Supplemental Figure S4). This corroborates earlier results obtained upon transient expression in cultured cells of Arabidopsis (Carrie et al., 2009). Furthermore, using GUS reporter lines, we showed that mFDX-like was expressed in roots, leaves and flowers. However, GUS activity was barely detectable in stem and siliques, and absent in cotyledons (Supplemental Figure S5). This expression pattern is compatible with a housekeeping mitochondrial function.

### The absence of mFDX-like does not affect plant growth

To gain more insights into the physiological role of mFDX-like, we first searched for knockout mutants from available T-DNA insertion lines. Three lines were tested: *mfdxl-1*: SALK_015333, *mfdxl-2*: WiscDsLoxHs204_01A and *mfdxl-3*: SAIL_37_H11 (Supplemental Figure S6A). We could obtain homozygous lines for *mfdxl-1* and *mfdxl-2* but western blots performed with an antibody raised against mFDX-like indicated that these two lines did not have neither a loss nor a decrease of mFDX-like when compared to wild-type (WT) plants (Supplemental Figure S6B-S6C). This was not so surprising as both T-DNA insertions are located in the 5’ UTR. Furthermore, we were unable to isolate homozygous lines for *mfdxl-3* as this mutant showed an arrest in the development of some seeds in the siliques of the heterozygous plants (Supplemental Figure S6D). Therefore, we tested the viability of the pollen grain using Alexander’s stain and observed that pollen-tetrads from the *mfdxl-3/-* plant indeed contained some dead cells (Supplemental Figure S6D). To test whether the lack of mFDX-like was responsible for such drastic phenotype, we generated two independent deletion lines using the CRISPR-Cas9 genome editing technology (Supplemental Figure S7). The mutants were backcrossed with WT plants to eliminate the Cas9 cassette and complemented by reintroducing the sequence coding for mFDX-like expressed under the constitutive 35S promoter in each line. The progeny from self-crossed plants was examined by western blot-based genotyping. Protein blot analysis on isolated mitochondria showed that the immune signal corresponding to the mature form of mFDX-like (∼ 15 kDa) was absent in both loss-of-function lines compared to WT plants (Figure 3A). Nevertheless, despite the absence of mFDX-like, none of these knock-out lines showed a growth phenotype when grown under standard conditions (Figure 3B). Additionally, this result discredited the potential lethality due to a lack of mFDX-like in the *mfdxl-3* T-DNA mutant line. Overall, this functional analysis demonstrated that mFDX-like is not essential for plant growth and development.

**Figure 3.**
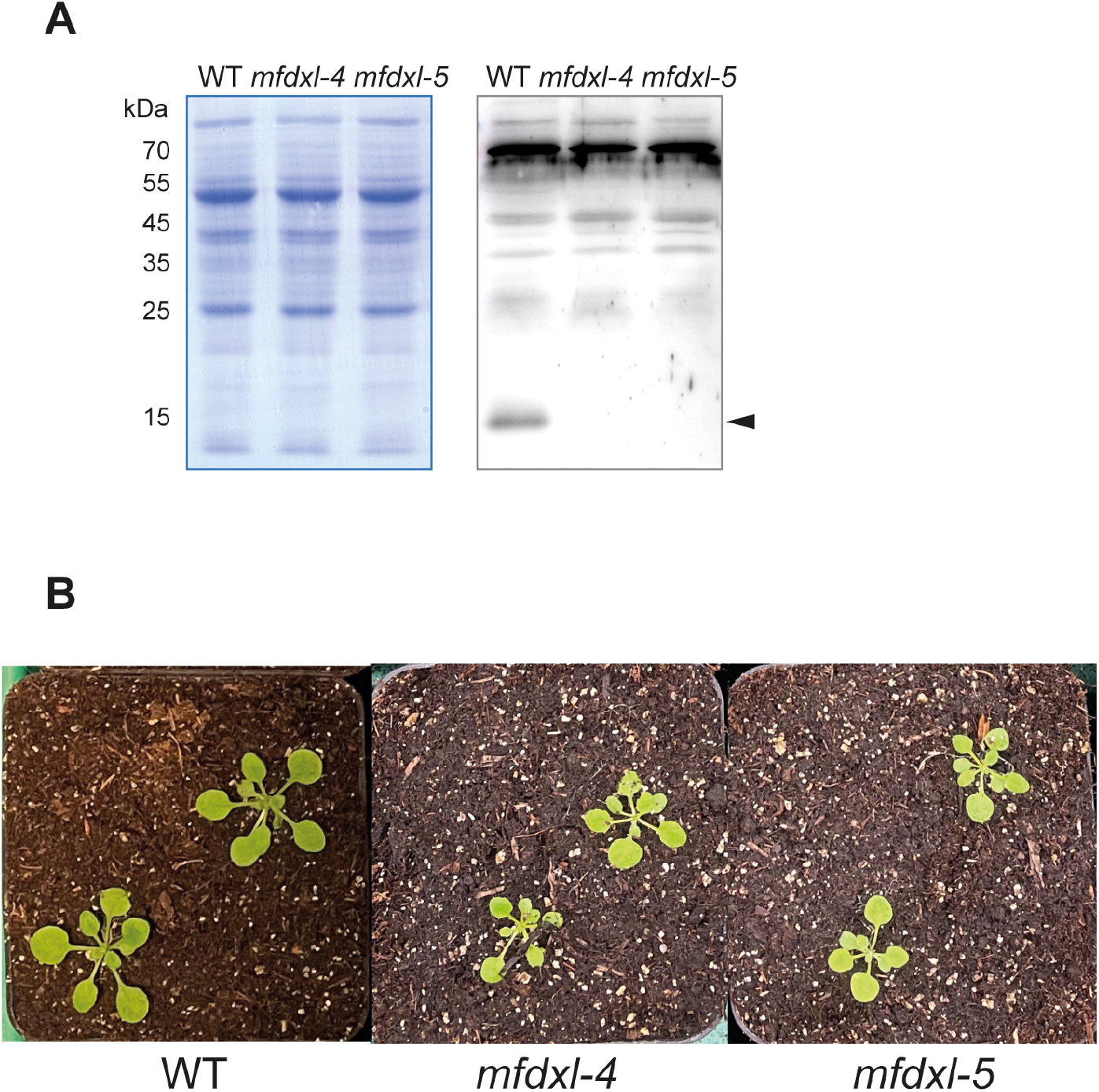
The absence of mFDX-like does not affect plant growth. Two independent deletion lines were generated using the CRISPR-Cas9 genome editing technology. A. Mitochondria were isolated from the lines and mitochondrial proteins were analyzed by SDS-PAGE followed by immunoblot experiments using an anti-mFDX serum. Left panel: Coomassie staining of the membrane. Right panel: Signals obtained after incubation of the membrane using the anti-mFDX-like antibodies. The position of the mature form of mFDX-like is indicated by an arrowhead. The aspecific signals observed in the top part of the blot serve as loading control. B. Representation images of 20-day-old seedlings grown on soil under standard growth conditions.

### mFDX-like is essential for supercomplex formation between complex I and complex III_2_

As already mentioned above, mFDX-like is associated with complex I (Hansen et al., 2018, Klusch et al., 2021). Consequently, we analyzed complex I levels in *mfdx-like* knock-out lines. To this end, mitochondria were isolated, complexes solubilized with the mild detergent digitonin and separated under native electrophoresis conditions. The gel was then stained with Coomassie blue to visualize the OXPHOS complexes. The bands corresponding to complex I, complex III dimer (III_2_) and complex V presented a similar intensity in the deletion lines in comparison with the WT and complemented lines (Figure 4). However, the bands corresponding to complex I-containing supercomplexes were not detectable in both knock-out lines (Figure 4). When mFDX-like was reintroduced, these supercomplexes were detectable again, demonstrating that mFDX-like is required for the formation of complex-I-containing supercomplexes in Arabidopsis.

**Figure 4.**
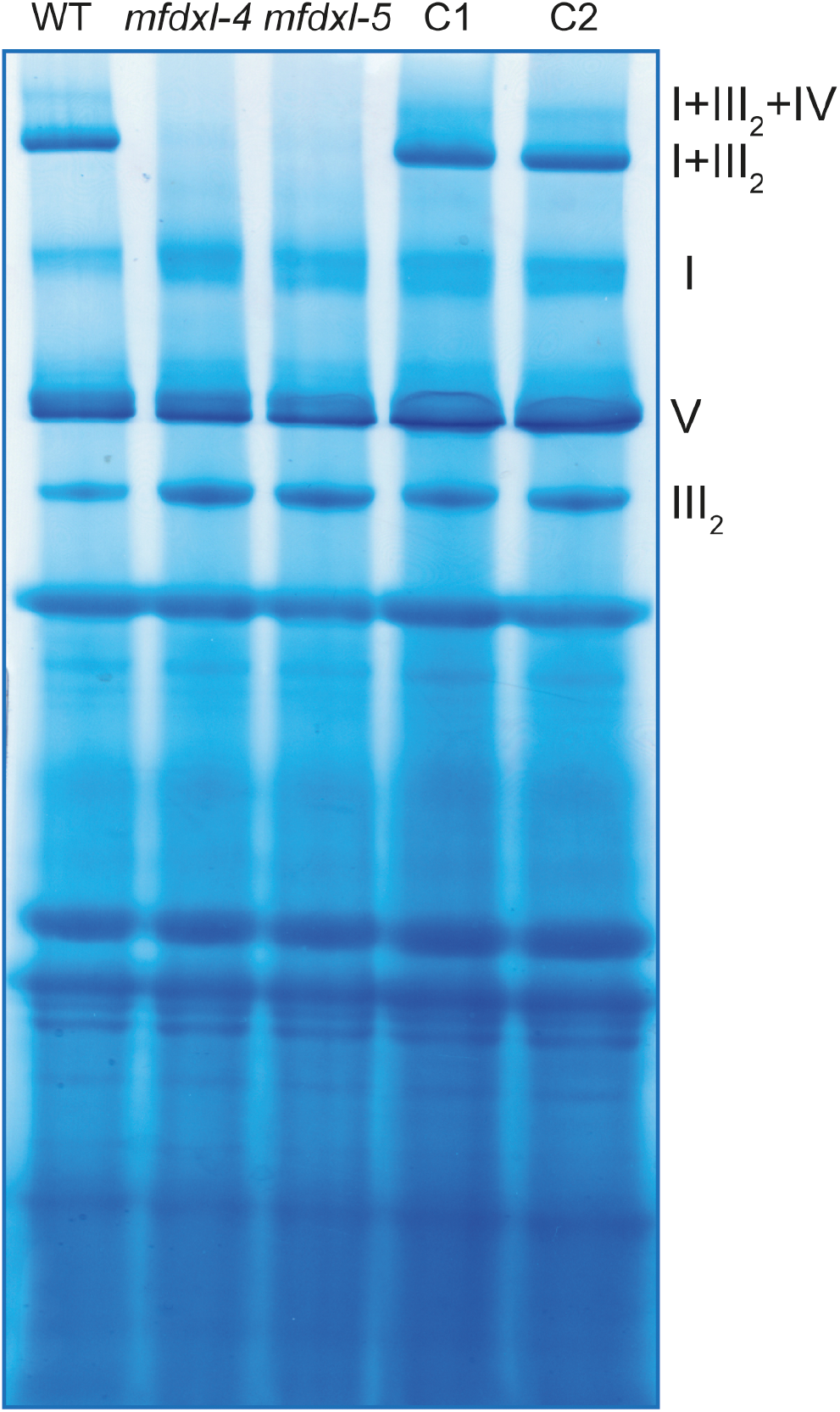
mFDX-like is essential for supercomplex formation. Blue native PAGE analysis of mitochondrial complexes. Mitochondria were solubilized using digitonin and the complexes separated on a native gel. After migration, the gel was stained with Coomassie blue. OXPHOS complexes I, III_2_ and V and supercomplexes (I+III_2_, I+III_2_+IV) are indicated on the right. C1 and C2 are the complemented lines of mutants *mfdxl-4* and *mfdxl-5* respectively.

### mFDX-like is important for the assembly of the membrane arm of complex I

To gain further insights into the role of mFDX-like in respiratory complex I formation, we performed additional native gels and transferred the multiprotein complexes on membranes (Figure 5A). Assembly intermediates and abundance of complex I were visualized by immunoblotting using antibodies recognizing specifically certain subunits or assembly factors of the complex I. As previously observed, no supercomplexes I+III_2_ were detected in Arabidopsis cas9-edited lines when antibodies raised against the complex I subunit carbonic anhydrase 2 (CA2) were used. This confirmed the importance of mFDX-like for the formation of complex-I-containing supercomplexes. Interestingly, the absence of mFDX-like led to the appearance of additional bands of a size both larger and smaller than mature complex I (Figure 5B). To determine whether the additional bands detected with the anti-CA2 antibodies represented assembly intermediates and not degradation products, we used anti-GLDH antibodies as GLDH is an assembly factor of complex I present in assembly intermediates but absent in mature complex I and its degradation products (Schimmeyer et al., 2016, Schertl et al., 2012, Ligas et al., 2019). The same band pattern was detected with the anti-CA2 or anti-GLDH antibodies in the deletion lines (Figure 5C). This indicated that these bands corresponded to assembly intermediates, and thus demonstrated that the assembly pathway of complex I is impaired in the absence of mFDX-like. Of note, the monomeric GLDH was not detected in the deletion lines while it was visible in both WT and complemented lines (Figure 5C; lower part of the native gel). The fact that the assembly intermediates were also slightly accumulated in the complementation lines compared with WT lines suggests that the complemented lines are not fully rescued.

**Figure 5.**
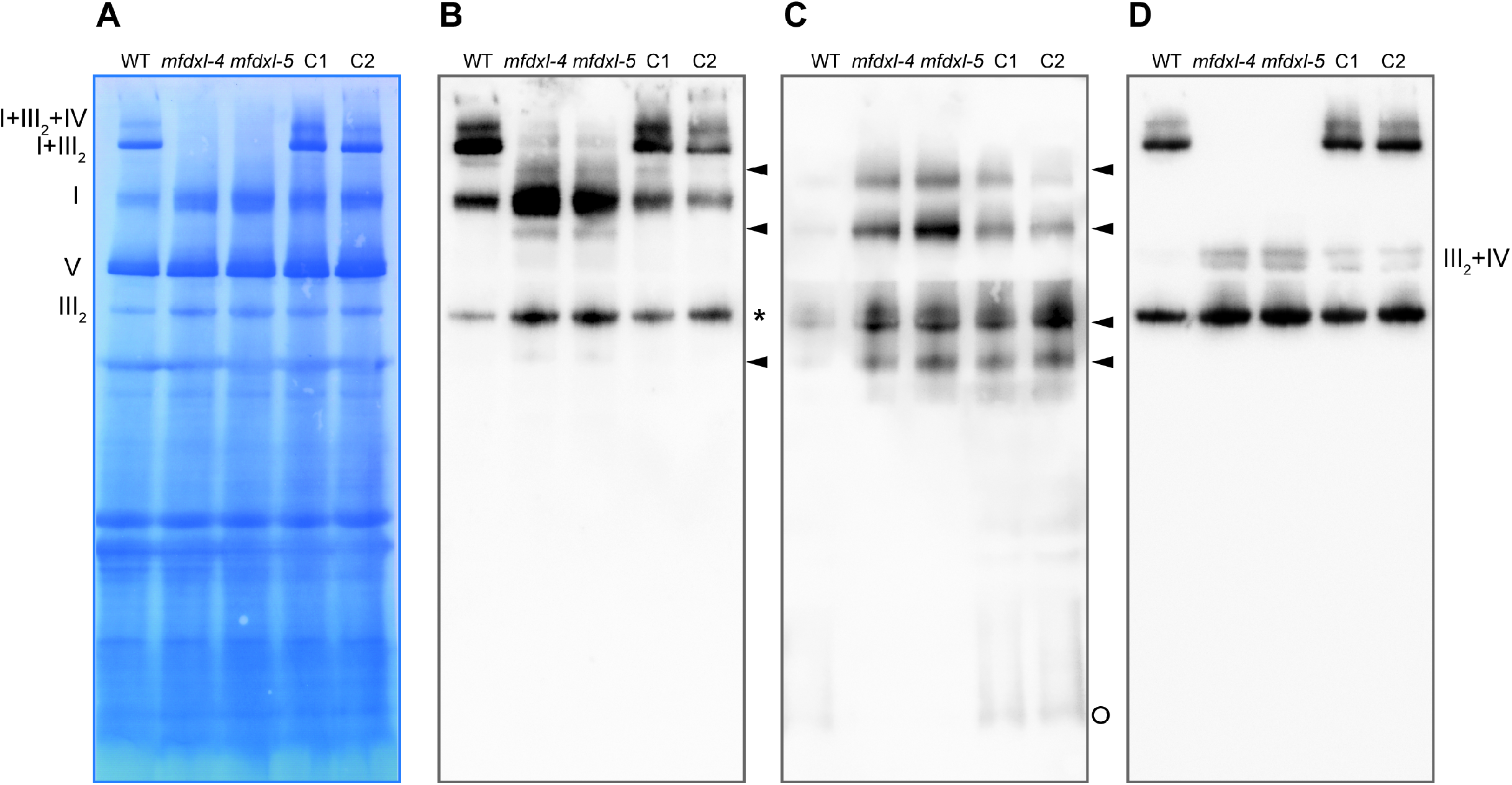
mFDX-like is important for the assembly of complex I. Blue native PAGE analysis of mitochondrial complexes. Mitochondria were solubilized using digitonin and the complexes separated on a native gel. After migration, the complexes were transferred on a PVDF membrane which was used for immunoblot analyses. C1 and C2 are the complemented lines of mutants *mfdxl-4* and *mfdxl-5* respectively. A. Coomassie staining of the membrane after transfer, the position of the OXPHOS complexes and supercomplexes is indicated on the left (I: complex I, III_2_: complex III dimer, V, complex V). B. Signals obtained after incubation of the membrane with the anti-CA2 antibodies. Complex I related complexes accumulated in the deletion mutants are indicated with an arrowhead. *: aspecific reaction of complex III_2_ with the ECL reagent. C. Signals obtained after incubation of the membrane with the anti-GLDH antibodies. Complex I assembly intermediates are indicated with an arrowhead. Free GLDH running close to the bottom of the gel is indicated with a circle. D. Signals obtained after incubation of the membrane with the anti-RISP antibodies. The additional signals running at the level of complex V likely represent the supercomplexes composed of complex III dimer and either form of complex IV (III_2_+IV). The same membrane has been used to perform the three immunoblot analyses.

Both anti-CA2 and anti-GLDH antibodies detect at least one band that is larger than complex I. Because of the presence of GLDH, this band likely contains the last and most stable assembly intermediate of complex I, complex I* (Ligas et al., 2019). To test if such band could correspond to an association of complex I* with complex III_2_ (the observed size of this band corresponds to the expected size for such an association), we performed an immunoblot detection with antibodies raised against a subunit of mitochondrial complex III_2_, the Rieske iron-sulfur protein (RISP). In these conditions, no complex I-containing supercomplexes were detected but instead an increase in the abundance of the supercomplexes composed of complex III_2_ and complex IV (Figure 5D). The largest GLDH-containing band is also not detected with the anti-RISP antibodies (Figure 5D), thus excluding that this band corresponds to a supercomplex I*+III_2_. In conclusion, the absence of mFDX-like impairs the assembly of complex I and leads to the accumulation of assembly intermediates such as complex I*.

### CA2 is important for the assembly of the P_D_ domain

Complex I* lacks the P_D_ module of the membrane arm. However, mFDX-like is part of the bridge domain linking the matrix arm and the CA domain that is attached to the P_P_ module of the membrane arm (Meyer et al., 2022); as such mFDX-like is not expected to directly interact with the P_D_ module. Therefore, the accumulation of complex I* in the deletion mutants cannot be explained by a direct role of mFDX-like in the assembly of the P_D_ module. One possible role of mFDX-like and the bridge domain is to facilitate the attachment of the P_D_ module on complex I* by triggering a conformational rearrangement of the membrane arm of complex I*. Such hypothesis is supported by the structures of complex I and complex I* in cauliflower, in which the CA domain is shifted by 6 Å between both structures (Soufari et al., 2020). Based on these data, a mutant impaired in the CA domain should also be impaired in the assembly of the P_D_ module. Because the CA domain is assembled early, no assembly intermediates can be detected in mutants lacking the CA subunits (Meyer et al., 2011). Nevertheless, assembly intermediates might be below the detection limit in such mutants as they accumulate very low levels of complex I (Perales et al., 2005, Fromm et al., 2016). To overcome this issue, we crossed *ndufs4*, a mutant accumulating the membrane arm, the P_P_ module and its assembly intermediates (Meyer et al., 2009, Kühn et al., 2015) with a *ca2* mutant in which the assembly of complex I is limited by the assembly of the CA domain. We performed reciprocal crosses to rule out a potential maternal effect, and obtained *ndufs4 ca2* and *ca2 ndufs4* lines. We then analyzed the assembly of complex I using native gel electrophoresis followed by immunodetection of Nad6, a subunit of the P_P_ module of complex I. In *ndufs4*, the full membrane arm can be assembled but in the double mutants, the membrane arm is not detectable, instead the P_P_ module accumulates (Figure 6). This indicates that the assembly is stalled, and that the P_D_ module cannot be efficiently associated with the P_P_ module in the absence of CA2. Therefore, both mFDX-like and CA2 are needed for the correct assembly of the P_P_ module.

**Figure 6.**
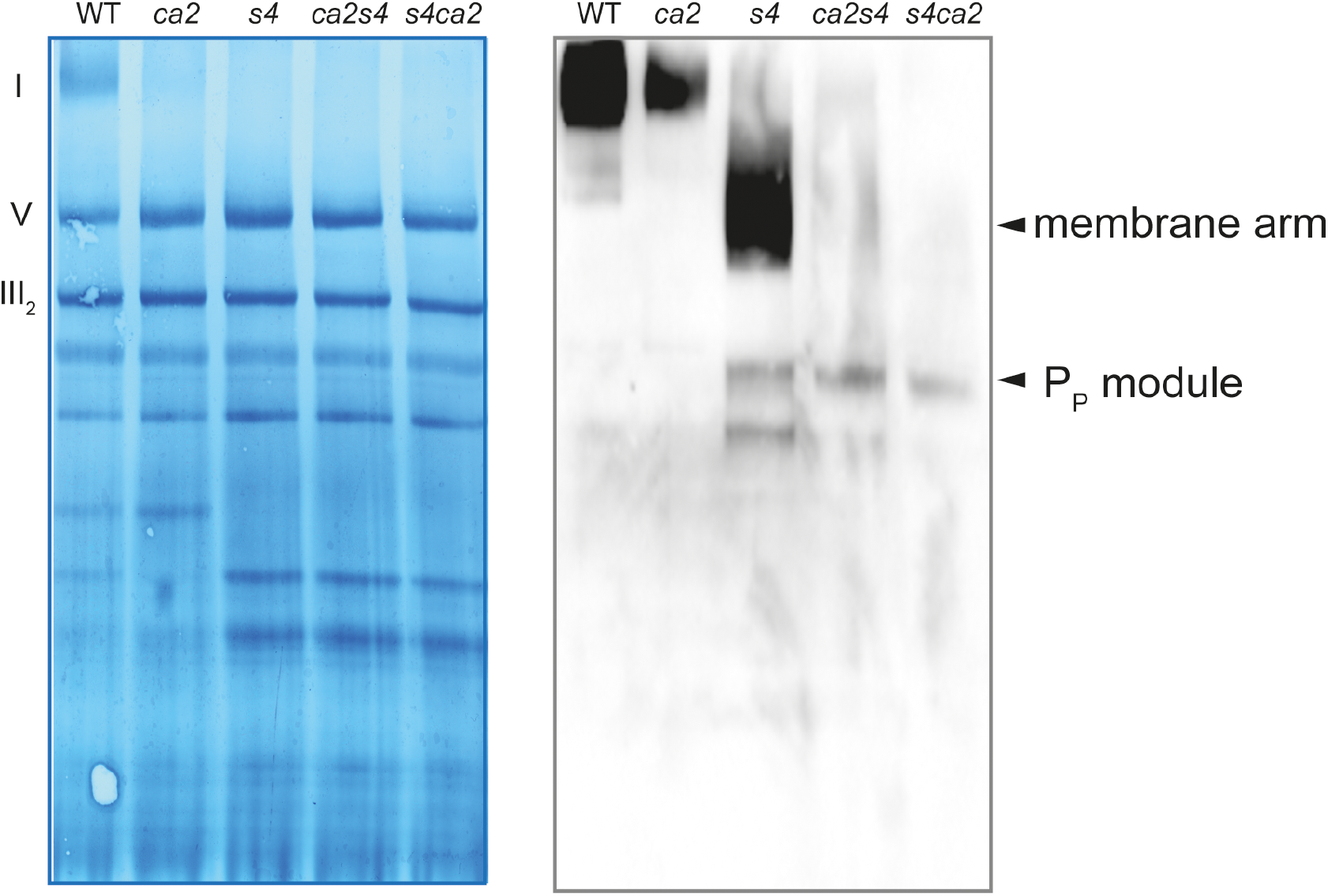
CA2 is important for the assembly of the P_D_ module. Blue native PAGE analysis of mitochondrial complexes. Mitochondria were solubilized using dodecyl-maltoside and the complexes separated on a native gel. After migration, the complexes were transferred on a PVDF membrane which was used for immunoblot analyses. Left panel: Coomassie staining of the membrane after transfer, the position of the OXPHOS complexes is indicated on the left (I: complex I, III_2_: complex III dimer, V, complex V). Right panel: Signals obtained after incubation of the membrane with the anti-Nad6 antibodies. Complex I assembly intermediates accumulating in the mutants are indicated on the right. *ca2*: mutant lacking the subunit CA2, *s4*: mutant lacking the subunit NDUFS4, *ca2s4* and *s4ca2*: double mutants lacking CA2 and NDUFS4.

### The role of mFDX-like in supercomplex formation is likely independent of NDUA11

The structure of supercomplexes has never been reported in plants so far. Based on the structure of supercomplexes I+III_2_ isolated from mammalian (Gu et al., 2016, Letts et al., 2019) or from Tetrahymena (Zhou et al., 2022), two main contact sites between complex I and complex III_2_ have been highlighted at the level of the membrane arm. The first one is formed by the NDUA11 subunit of complex I and the UQCRB (QCR7) subunit of complex III_2_ whereas the second one occurs between NDUB4 and NDUB9 subunits of complex I and UQCRC1 (MPPbeta) subunit of complex III_2_ (Letts et al., 2019). NDUA11 is not present in the structure of Arabidopsis complex I (Soufari et al., 2020) but if its binding site is conserved between opisthokonts and plants, NDUA11 should be located just underneath mFDX-like. Hence, a possible role of mFDX-like could be to stabilize NDUA11. If this is true, NDUA11 would be unable to mediate the interaction with complex III_2_ in the absence of mFDX-like, leading to the absence of the supercomplex I+III_2_. To test this hypothesis, we isolated a homozygous T-DNA insertion line in the *NDUA11* gene. In these plants, NDUA11 gene product was not detected from isolated mitochondria by immunoblot assay using specific antibodies (Michaud et al., 2016) (Figure 7A). Yet, knock-out Arabidopsis *ndua11* showed normal growth and development when compared to WT plants (Figure 7B). Despite the absence of a macroscopic phenotype, we checked whether a *ndua11* mutant could be affected in OXPHOS complexes using native gel electrophoresis coupled with immunoblot analyses. Similar to what was observed in a *mfdx-like* mutant, complex I-containing supercomplexes were undetectable in the absence of NDUA11 (Figure 7C), confirming that NDUA11 is essential for the interaction between complex I and complex III_2_ within the supercomplex. We next tested if the absence of mFDX-like could disturb the recruitment of NDUA11 in the respiratory complex I. For this purpose, immunoblot analysis with anti-NDUA11 antibodies was assayed on separated OXPHOS complexes from WT and *mfdx-like* plants. In all genetic backgrounds, NDUA11 associated with complex I (Figure 7D). Therefore, the absence of supercomplexes I+III_2_ in *mfdx-like* knock-out lines does not result from the absence of NDUA11.

**Figure 7.**
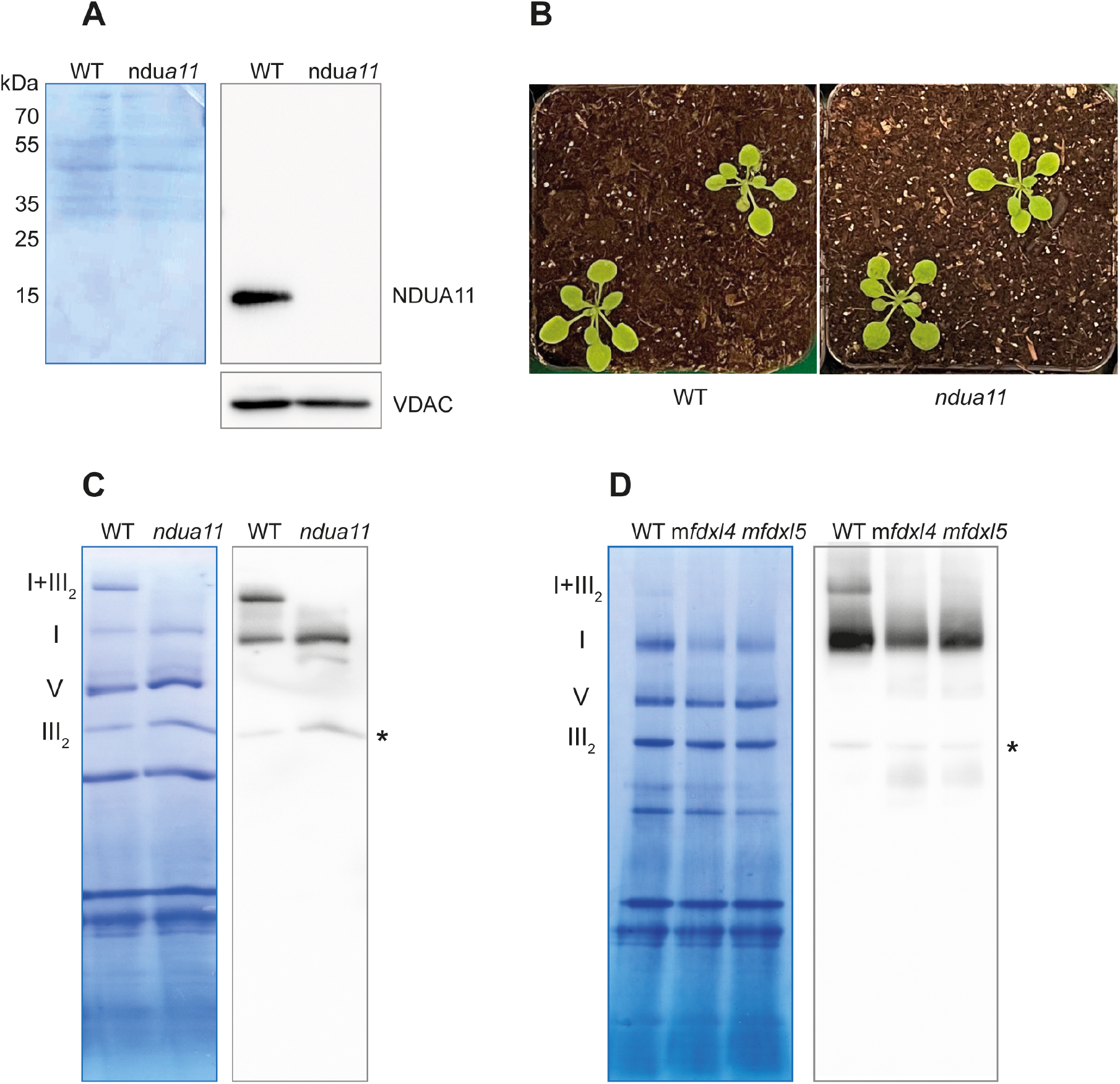
NDUFA11 is essential for supercomplex formation but its association with complex I is not affected in m*fdxl* mutants. A. Mitochondria were isolated from the *ndua11* and WT plants and mitochondrial proteins were separated by SDS-PAGE and analyzed by immunoblot analyses. Left panel: Coomassie staining of the membrane. Right panels: Signals obtained after incubation of the membrane with first the anti-NDUA11 antibodies and second the anti-VDAC antibody. The detection of VDAC is used as a loading control. B. Image of 20-day-old seedlings grown on soil under standard growth conditions. C. Blue native PAGE analysis of mitochondrial complexes. Mitochondria were solubilized using digitonin and the complexes separated on a native gel. After migration, the complexes were transferred on a PVDF membrane which was used for immunoblot analyses. left panel: Coomassie staining of the membrane after the transfer, the position of the OXPHOS complexes is indicated on the left (I+III_2_: supercomplex I+II_I2_, I: complex I, III_2_: complex III dimer, V, complex V). right panel: Signals obtained after incubation of the membrane with the anti-CA2 antibodies. *: aspecific reaction of complex III_2_ with the ECL reagent. D. Blue native PAGE analysis of mitochondrial complexes. Mitochondria were solubilized using digitonin and the complexes separated on a native gel. After migration, the complexes were transferred on a PVDF membrane which was used for immunoblot analyses. left panel: Coomassie staining of the membrane after the transfer, the position of the OXPHOS complexes is indicated on the left (I+III_2_: supercomplex I+III_2_, I: complex I, III_2_: complex III dimer, V, complex V). right panel: Signals obtained after incubation of the membrane with the anti-NDUA11 antibodies. *: aspecific reaction of complex III_2_ with the ECL reagent.

## Discussion

### Evolution of the function of mFDX-like

According to our phylogenetic analysis, Arabidopsis FDXs from eukaryotic organisms are grouped into three clades: the plastidial FDXs, the classical mitochondrial FDXs and a clade containing both classical and divergent mitochondrial FDXs (based on the presence or the absence of one or more of the cysteines required for the coordination of the Fe-S cluster) that includes AtmFDX-like and the two other FDXs previously shown to be associated with complex I. Interestingly, additional orthologs are present in this branch and represent potential candidates for complex I-associated FDXs. However, this would have to be experimentally validated. Also, this clade contains proteins from organisms belonging to distinct eukaryotic groups (Supplemental Table S1), suggesting that the association of an FDX with complex I is an ancestral feature of this complex.

The *in vitro* characterization of Arabidopsis mFDX-like demonstrated that it does not bind an Fe-S cluster according to the lack of a cysteine ligand. Hence, mFDX-like does not behave as a classical FDX, and its involvement in electron transfer reactions is unlikely. The absence of cysteine ligands in most algal orthologs suggests that these residues do not have an important structural or functional role. Still, because the cysteines have been conserved during evolution in mFDX-like from terrestrial plants, it would be interesting to test whether they might be important for the role of mFDX-like in these organisms, for instance by making functional complementation assays. Altogether these phylogenetic and biochemical data suggest that the function of mFDX-like has likely evolved from a conventional enzymatic electron transfer function towards a strictly structural one.

It is worth mentioning that mFDX-like has never been identified as a complex I subunit in proteomic studies in Arabidopsis. This likely results from the consequence of the use of native gel electrophoresis as the bridge domain was only observed so far in sample preparations that do not include gel electrophoresis. Based on the analysis of the number of protein copies in a single mitochondrion, mFDX-like seems to be a quite abundant protein, being present at levels globally similar to other complex I subunits and much more abundant than the two regular mFDX1/2 (Fuchs et al., 2020). However, it remains to be determined whether mFDX-like is exclusively associated with respiratory complex I or if it can also play independent, non-essential roles as a soluble protein in the mitochondrial matrix.

### Assembly of complex I

In the mFDX-like knock-out lines, no alteration of complex I levels have been observed but several assembly intermediates accumulate (Figure 5). We used two different antibodies to detect these intermediates, the anti-CA2 and anti-GLDH antibodies. CA2 is a subunit of the CA domain that is assembled early and is attached on the P_P_ module of the membrane arm of complex I. GLDH is an assembly factor that is associated with three assembly intermediates including the P_P_ module and complex I* that is formed by the attachment of the almost fully assembled matrix arm on the P_P_ module (Ligas et al., 2019). The accumulation of complex I* in the mFDX-like deletion lines indicates that the final step of the assembly is impaired. Thus, mFDX-like should be associated with complex I* to proceed to the next step of the assembly. This step is the binding of the distal part of the membrane arm, the P_D_ module. For this to occur, GLDH must be removed, the P_D_ module attached to complex I* and the NDUP1 subunit be bound across the P_D_ and P_P_ modules at the position where GLDH was bound (Meyer et al., 2022). We observed an absence of free GLDH in the mFDX-like deletion lines, indicating that the bridge domain is important for the release of GLDH from complex I assembly intermediates. However, as complex I can be fully assembled in the mFDX-like deletion lines, the release of GLDH can occur in absence of mFDX-like but this step appears less efficient in the deletion lines than in WT plants. Our data demonstrate that mFDX-like is important for the transition from complex I* to complex I.

The analysis of the two reciprocal double mutants lacking NDUFS4 and CA2 also points towards a role of CA2 is the assembly of the P_D_ module and suggests that mFDX-like and the CA domain are important for the efficient assembly of the P_D_ module. The structural characterization of complex I and complex I* in cauliflower supports a common function of mFDX-like and the CA domain in the transition from complex I* to complex I. The position of the CA domain was found to be shifted by 6 Å towards the missing P_D_ module between complex I and complex I* (Soufari et al., 2020). A possible mechanism integrating all these results would be that upon building of the bridge domain (binding of mFDX-like), the CA domain is pushed away by 6 Å, allowing the release of GLDH from complex I* and the binding of the P_D_ domain.

In *ca2*, the CA domain is present and looks similar than in WT (Sunderhaus et al., 2006), indicating that CA2 is replaced by another carbonic anhydrase subunit. This might affect the position of the CA domain on the membrane as the N-terminal α-helix of CA2 is involved in the interaction between the CA domain and the P_P_ module (Soufari et al., 2020, Klusch et al., 2021, Maldonado et al., 2020). Low levels of complex I and supercomplex I+III_2_ can be assembled in *ca2* (Perales et al., 2005, Fromm et al., 2016), indicating that the unusual CA domain in *ca2* remains limiting for the assembly but the assembly process is not disturbed as no assembly intermediates accumulate. However, in the *ndufs4 ca2* double mutants, the binding of the P_D_ module on the P_P_ module is impaired (Figure 6). The main difference between *ca2* and the *ndufs4 ca2* mutants is the presence or absence of the matrix arm. This suggests that, when the CA domain does not contain CA2, the matrix arm should be attached to the P_P_ domain to allow the binding of the P_D_ domain. Whether or not this is linked to the formation of the bridge domain between the matrix arm and the CA domain remains to be explored.

In summary, both the bridge domain including mFDX-like and the CA domain are important for complex I assembly as the bridge domain likely influences the positioning of the CA domain on the P_P_ module, the release of GLDH and the binding of the P_D_ domain. However, mFDX-like is not directly responsible for the release of GLDH as GLDH removal and complex I assembly can occur in the deletion lines, albeit inefficiently. However, the complex I accumulating in these lines should have a different conformation than the one in the WT line, with the P_D_ module being misoriented. Unfortunately, this cannot be currently tested as structural work in Arabidopsis has, to date, only be successful using cell cultures as starting material.

### Formation of supercomplexes

Based on the observation that complex I-containing supercomplexes are undetectable in the mFDX-like deletion lines, we concluded that the complex I that accumulates is unable to associate with complex III_2_. This suggests either that mFDX-like is involved in the interaction with complex III_2_ or that it is important for the formation of one of the contact points between both complexes. No structures of supercomplex I+III_2_ from plants are available. Yet, tomography imaging revealed that the angle between complex I and complex III_2_ is larger in plants than in opisthokonts (Davies et al., 2018). As the bridge domain of complex I faces complex III_2_, this could explain why more space is required between both complexes. Recently, the structure of the supercomplex I+III_2_ from *Tetrahymena* has been solved (Zhou et al., 2022). The bridge domain of complex I also contains a FDX, but also additional proteins that interact with complex III_2_ subunits. Hence, it is possible that a similar organization exists in plants as well and that additional uncharacterized proteins present in the bridge domain and in the supercomplex I+III_2_ remain to be identified in plants.

Two main contact points have been described at the level of the membrane arm. First, the complex I NDUA11 subunit interacts with UQCRB (QCR7) from complex III_2_. In the Tetrahymena supercomplex I+III_2_ structure, these two subunits coordinate a zinc atom together (Zhou et al., 2022). The second contact point is between NDUB4 and NDUB9 subunits of the P_D_ module and UQCRC1 (MPP-β) from complex III_2_ (Letts et al., 2019). We have shown that NDUA11 is essential for supercomplex formation but that NDUA11 is present in the complex I accumulating in the mFDX-like deletion lines. In addition, our data suggest that the assembly of the P_D_ module is dependent on structural rearrangements triggered by mFDX-like. Altogether, these results point towards a structural role for mFDX-like to correctly position either NDUA11 or the P_D_ module. In the absence of mFDX-like, the P_D_ domain can still associate with complex I* but its position would be shifted, impairing the association with complex III_2_. As NDUA11 interacts with an α-helix of the subunit Nad5 in the P_D_ module, an inadequate position of the P_D_ module might influence the positioning of NDUA11 and also disturb the interaction with complex III_2_. Overall, we propose that mFDX-like and the bridge domain are essential for the correct organization of the membrane arm of complex I. To test further this hypothesis, the determination of the structure of complex I in the mFDX-like deletion lines would be required.

## Conclusion

We previously identified mFDX-like through a bioinformatic screen aimed at identifying genes encoding mitochondrial proteins involved in complex I proteostasis (Hansen et al., 2018). More recently, structural analyses associated mFDX-like proteins from Arabidopsis and the green alga Polytomella with the respiratory complex I (Klusch et al., 2021), reinforcing our predictions and demonstrating the robustness of our initial bioinformatic approach. In this work, we provide new insights into the biochemical properties and importance of mitochondrial FDX-like in the assembly pathway and supercomplex formation of the plant respiratory complex I. This suggests a key role for mFDX-like in the conformation of the membrane domain. Interestingly, the fact that the absence of mFDX-like had no influence on plant growth under standard growth conditions supports the idea that complex I-containing supercomplexes are dispensable for plant growth. Yet, future studies should aim at challenging the physiological function of respiratory supercomplexes in plants under stress, and several of the mutant lines we have created would thus represent valuable tools for these investigations.

## Material & Methods

### Sequence alignments and phylogenetic analysis

Arabidopsis mFDX protein sequences were retrieved from the Arabidopsis information resource (TAIR). Yeast and human mFDX amino acid sequences were collected from the Uniprot database (release 2022_03) whereas the complete protein sequences corresponding to *A. thaliana* mFDX-like orthologs were obtained using AtmFDX-like as a reference sequence and BLASTP tools at the Phytozome (v13) database. To retrieve the FDX sequences in selected eukaryotes (Supplemental Table S1), PSI-Blast searches were performed using mFDX1, mFDX2 or mFDX-like as bait. Phylogenetic and molecular evolutionary analyses were conducted using MEGA version 11 (Tamura et al., 2021).

Multiple sequence alignments were performed with the multiple sequence comparison by log-expectation (MUSCLE) program using default parameters. The phylogenetic tree was constructed with the minimum-evolution option using default parameters. Final graphic views of the aligned sequences were generated with the BioEdit software (version 7.2.5).

### Expression of proteins and *in vitro* biochemical assays

The sequences encoding the presumed mature forms of mFDX-like (AA 37-159), mFDX1 (AA 70-197), mFDX2 (AA 71-197) proteins were amplified from cDNA prepared from *A. thaliana* Col-0 leaf total RNA. Primers used are listed in Supplemental Table S2. The PCR products were cloned into pET15b expression vector (Novagen) between NdeI and BamHI restriction sites, resulting in the introduction of a hexa-His tag at the N-terminus of the expressed proteins. Expression of proteins in the *E. coli* BL21(DE3) pSBET strain transformed with recombinant pET15b plasmids was induced at exponential phase with 100 μM of isopropyl β-D-1-thiogalactopyranoside during 4h at 37°C in 2.4 L cultures. Bacteria were then collected by centrifugation at 6,000 g during 20 min and resuspended in 30 mM Tris-HCl pH 8.0, 200 mM NaCl, 10 mM imidazole lysis buffer and eventually conserved at - 20°C. Cell lysis was completed by three cycles of 1 min sonication on ice and the soluble fraction was separated from insoluble fractions and cellular debris by centrifugation at 40,000 g during 40 min. The soluble fractions were loaded on a Ni-NTA (Ni^2+^-nitrilotriacetate)– agarose resin equilibrated with a 30 mM Tris-HCl pH 8.0, 200 mM NaCl, 10 mM imidazole buffer. After a washing step, elution of captured His-tagged proteins from the column is accomplished by using the same buffer containing 250 mM imidazole. The fractions containing the proteins of interest were pooled, concentrated and imidazole was removed by repeated cycles of ultrafiltration using a YM10 membrane in an Amicon cell under nitrogen pressure. Purity and integrity of purified recombinant proteins were analysed by 15% SDS/PAGE. Protein concentrations were determined by measuring the absorbance at 280 nm and using theoretical molar absorption coefficients of 2980 M^-1^.cm-^1^ for AtmFDX1 and AtmFDX2 and of 15470 M^-1^.cm^-1^ for AtmFDX-like and AtmFDX-like L85C. The proteins were stored at −20°C.

### Confocal laser microscopy imaging

Fragment comprising 1,000-bp region located upstream of the ATG translation start codon of *mFDX-like* gene to last codon before stop codon were amplified from wild-type genomic DNA using primers listed in Supplemental Table S2 and cloned into pGWB4 vector (Invitrogen). After Agrobacterium mediated plant transformation, progenies were selected on ½ MS plates containing 0.7% (w/v) phytoagar, 50 μg/mL hygromycin antibiotic, and then cultivated in soil. Two-week-old roots grown on a ½ MS (0.8% agarose) media were vacuum infiltrated with 100 nM of Mitotracker Orange CMTMRos (ThermoFischer, M7510), diluted in ½ MS, for 5 min and incubated 15 min in darkness. Fluorescence of the mFDX-like-GFP fusion protein was observed on a confocal microscope (Zeiss, LSM780) and a Zeiss Zen software. Signals were detected according to the following excitation/emission wavelengths: GFP (488 nm/495-535 nm) and Mitotracker (561 nm/575-630 nm). Pictures were analysed using the ImageJ software (https://imagej.nih.gov/ij/) where only contrast and brightness were adjusted.

### Histochemical GUS staining

Full-length genomic sequence from the promoter region 1,000-bp upstream of the ATG translation start codon of *mFDX-like* gene to last codon before stop codon were amplified from genomic DNA of WT plants and cloned into pGWB3 vector (Invitrogen). Primers used are listed in Supplemental Table S2. Transgenic plants were selected on ½ MS plates supplemented with 50 μg/mL hygromycin and then transferred in soil after 7 days. For histochemical staining, tissues from transformed T2 lines were incubated in GUS staining solution composed of 50 mM NaH_2_PO_4_ pH 7.0, 10 mM EDTA pH 8.0, 0.2% (v/v) Triton X-100, 2 mM of each hexacyanoferrate(III/II) (ferricyanide/ferrocyanide) and 2 mM X-gluc (5-bromo-4-chloro-3-indolylβ -D-glucuronide cyclohexylamine), followed by vacuum infiltration (3 cycles of 15 min). After overnight staining at 37°C, reaction was stopped with two washes in water and samples were placed in 70% (v/v) ethanol. The stained plants were observed and photographed under a Leica MZ9.5 stereo microscope. Three independent transgenic lines were analysed for expression of the GUS reporter that each showed similar expression patterns.

### Obtention of plant lines

The T-DNA lines SALK_015333, SAIL_434_E06, and WiscDsLoxHs204_01A were obtained from the Nottingham Arabidopsis Stock Center and homozygous seedlings were obtained after selfing and confirmed by genotyping. CRISPR-Cas9 was used to create deletions in the gene encoding mFDX-like (Supplemental Figure S7A). First, to generate the PCR template pJF1033, the two sgRNA encoding cassettes were amplified from plasmid pHEE2E-TRI (Wang et al., 2015) with primers oJF212 and oJF215, the PCR product digested with HindIII and SpeI and ligated into equally cut pGGZ001 (Lampropoulos et al., 2013). The pair of guide RNAs was cloned in the pJF1031 vector (Ruf et al., 2019) that allows the expression of Cas9 under an egg-cell-specific promoter. First PCR primers containing the guide RNAs (oJF398 and oJF399) were used to amplify a fragment of the pJF1033 plasmid and the PCR product was inserted into the BsaI sites of pJF1031. The resulting binary plasmid, pJF1120 (protospacer sequences AAACCGGAAAAGTGAGTGA & TGCGAGGTTCAGATCGCAG), is identical to pHEE2E-TRI except for the protospacer parts of the guide RNAs. pJF1120 was used to transform *Agrobacterium tumefaciens* GV3101 and the resulting bacteria were used to transform WT Col-0 plants by floral dip. Seeds of the T0 plants were screened on hygromycin. Resistant T1 plants were screened by PCR to detect deletion in the mFDX-like gene using primers mFDXl_ScrF and mFDXl_ScrR. Heterozygous plants were selfed and their progeny screened to obtain homozygous plants in which the wild type allele is not detectable anymore by PCR (Supplemental Figure S7B). Sequencing of the PCR products obtained from the mutant plants identified two independent deletions (Supplemental Figure S7C). Both lines were backcrossed with WT plants to remove the Cas9 cassette and homozygous deletion mutants were screened by PCR. The elimination of the Cas9 cassette was confirmed by PCR using primers to detect Cas9 (Cas9_F and Cas9_R) and by testing the sensitivity to hygromycin of the back crossed homozygous lines. The two lines were named *mfdxl-4* and *mfdxl-5*. For the complementation, the genomic region corresponding to the coding sequence of mFDX-like was amplified by PCR using primers mFDXl_ComF and mFDXl_ComR and cloned into the pGWB2 vector. The resulting vector was transformed into *Agrobacterium tumefaciens* GV3101 and this strain was used to transform the two deletion lines by floral dip. Transformants were screened on hygromycin. A hygromycin sensitivity test was performed on T3 plants to identify complemented lines homozygous for the complementation construct. The sequences of all the primers used are given in Supplemental Table S2.

### Plant cultivation

Seeds were surface sterilized with ethanol 70% (v/v) containing 0.05% (v/v) tween20 and plated on MS media containing 1% (w/v) sucrose. Plates were incubated under long day conditions (16h light 120 μE, 22°C and 8h darkness, 20°C, the humidity was not controlled and oscillated between 40 and 65%). 10-day-old seedlings were transferred on soil and grown under the same long day conditions.

### Pollen viability assay

Inflorescences were collected from 6-week-old plants and fixed in a 6 ethanol:3 chloroform: 1 glacial acetic acid) during 3h. After drying, flowers or buds were dissected, place on microscope slide and anthers were stained with a modified Alexander’s stain composed of 9.5% (v/v) ethanol, 0.01% (w/v) malachite green, 25% (v/v) glycerol, 0.05% (w/v) acid fuchsin, 0.005% (w/v) Orange G and 4% (v/v) glacial acetic acid (Peterson et al., 2010). Preparations were quickly heated and observed under a light microscope (Zeiss Axio Observer.A1m equipped with a HBO 100 camera).

### Mitochondrion isolation

Mitochondria were isolated from 4-week-old plants according to (Meyer et al., 2009). Briefly, the aerial parts of the plant were ground in extraction buffer (0.3 M sucrose, 15 mM potassium pyrophosphate, 2 mM EDTA, 10 mM KH_2_PO_4_, 1% (w/v) polyvinylpyrollidone-40, 1% (w/v) bovine serum albumin, 20 mM sodium ascorbate, pH 7.5) in a cold mortar. After filtration, two rounds of differential centrifugations (2.000 g/20.000 g) were performed to obtain a mitochondria-enriched pellet. After resuspension in wash buffer (0.3 M sucrose, 1 mM EGTA, 10 mM MOPS/KOH pH 7.2), the fraction was loaded on a discontinuous Percoll gradient comprised of three layers (18%-25%-50%) which was centrifuged at 40.000 g for 45 min. The mitochondria were collected at the interface between the 25% and 50% layers and the Percoll was eliminated by performing several washes in wash buffer. Protein concentration was estimated using the Bradford assay (Rotiquant, Roth).

### Gel electrophoresis and blotting

For the SDS-PAGE analysis, 10 μg of mitochondrial proteins were loading on a 12% acrylamide gel. After migration, the proteins were transferred on a PVDF membrane (Immobilon-P, Millipore) using a wet blotting system. For the BN-PAGE, 100 μg of mitochondrial proteins were solubilized with either 5% (w/v) digitonin or 1% (w/v) β-D-dodecyl-maltoside and solubilized protein complexes were separated on a BN gel according to (Eubel et al., 2005). For some experiments, protein solubilization was performed directly after the isolation of mitochondria whereas for other experiments, mitochondria were stored frozen between isolation and solubilization. This storage step damages the supercomplexes and lower amounts of supercomplexes are detectable (Figure 6 and 7). After migration, the complexes were transferred on PVDF membrane (Immobilon-P, Millipore) using a wet blotting system. After transfer, the membranes were stained for 5 min in staining buffer (30% (v/v) methanol, 7% (v/v) acetic acid, 0.005% (w/v) Coomassie Blue R250). The background was reduced by washing the membranes in 30% (v/v) methanol, 7% (v/v) acetic acid. The stained membrane was scanned and the staining was fully removed using 100 % (v/v) methanol.

### Antibody production and western blot

Polyclonal antibodies were produced in rabbits against the purified mFDX-like (Agrobio, France) or the peptide RALRKKYARKDEDY specific to CA2 (Biogenes, Germany). The final bleeds were used for western blots. For western blot experiments, membranes were first blocked using a 5% milk solution and incubated with the primary antibodies for 16h at 4°C. Following three washes with TBS-T (20 mM Tris-HCl pH 7.4, 150 mM NaCl, 0.1 % (v/v) Tween 20), the membranes were incubated with the secondary antibodies conjugated with HRP for 1h at 4°C and again washed three times with TBS-T. Final detection was performed using the ECL prime detection reagent (Cytiva Amersham) and a CCD camera (Fusion FX, Vilber). The primary antibodies were used at the following dilutions: anti-mFDX-like 1/5000, anti-CA2 1/10000, anti-GLDH (Agrisera AS06182) 1/5000, anti-GDC (Agrisera AS05074) 1/1000, anti-RISP (Carrie et al., 2010) 1/10000, anti-Nad6 (Koprivova et al., 2010) 1/5000 and anti-NDUA11 (Michaud et al., 2016) 1/20000 and anti-VDAC (Blake et al., 2007) 1/10000.

## Accession numbers

Accession numbers are as follows: AT3G07480: mFDX-like, AT4G05450: mFDX1, AT4G21090: mFDX2, At1g47260: CA2, At5g67590: NDUS4, At2g42210: NDUA11.

## Acknowlegments

The work was financially supported by a grant of the Deutsche Forschungsgemeinschaft (ME 4174/3-1) to EHM; by the Kempe Foundations (Gunnar Öquist Fellowship) and TC4F to OK; by a funding from the Lorraine University of Excellence (LUE) to NR. We would like to thank Tiphaine Dhalleine (IAM, Univ. Lorraine) for the help with the purifications of mFDX-like proteins and Michael Tillich for the help in designing the guide RNAs.

## Author Contributions

Project design: OK, NR, EHM; Experimental work: HR, JPT, JF, CB, NR, EHM; Data analysis: HR, JPT, OK, NR, EHM; Paper writing: EHM with contributions from all authors.

## Supplemental information

**Supplemental Figure S1**. Amino acid sequence alignment of AtmFDX-like with mitochondrial ferredoxin homologues.

**Supplemental Figure S2**. Amino acid sequence alignment of selected plant and algal mFDX-like proteins.

**Supplemental Figure S3**. Purification steps and purity degree of recombinant proteins.

**Supplemental Figure S4**. Subcellular localization of AtmFDX-like in Arabidopsis.

**Supplemental Figure S5**. Histochemical localization of GUS in transgenic *Arabidopsis thaliana* plants containing the *pmFDX-like:mFDX-like:GUS*.

**Supplemental Figure S6**. Genetic analysis and characterization of *mfdx-like* T-DNA insertion mutants from *Arabidopsis thaliana*.

**Supplemental Figure S7**. CRISPR/Cas9 strategy to obtain deletion mutants

**Supplemental Table S1**. UniProt accession numbers of the proteins used for the phylogenetic study

**Supplemental Table S2**. Primers used in this study

