## Supplemental Data for "The mitochondrial ferredoxin-like is essential for the formation of complex I-containing respiratory supercomplexes in *Arabidopsis thaliana*"

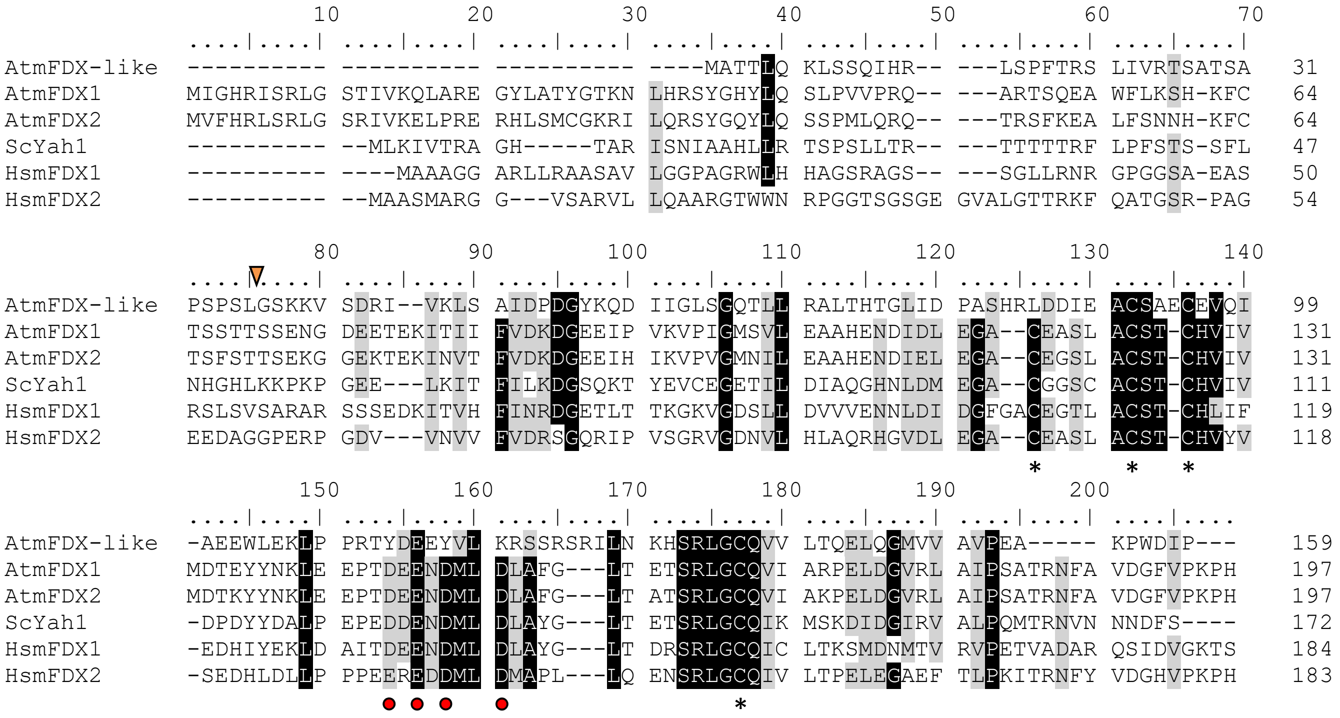


**Supplemental Figure S1. Amino acid sequence alignment of AtmFDX-like and mitochondrial ferredoxin homologues.** Conserved cysteine residues required for the [2Fe-2S] cluster coordination are pointed by asterisks whereas acidic amino acid residues a priori involved in the interaction with the ferredoxin reductase based on the structure of the adrenodoxin/adrenodoxin reductase complex (Muller et al., 2001) are indicated by red rings. Chosen cleavage sites for recombinant protein production are represented by an orange triangle. *A. thaliana* (At), *S. cerevisiae* (Sc) and *H. sapiens* (Hs). Shaded areas indicate at least 80% amino acid identity (black) or similarity (grey). The UniProt accession numbers are: AtmFDX-like: Q9SRR8, AtMFDX1: Q9M0V0, AtmFDX2: Q8S904, ScYah1: Q12184, HsmFDX1: P10109, HsmFDX2: Q6P4F2.


MULLER, J. J., LAPKO, A., BOURENKOV, G., RUCKPAUL, K. & HEINEMANN, U. 2001. Adrenodoxin reductase-adrenodoxin complex structure suggests electron transfer path in steroid biosynthesis. *J Biol Chem,* 276**,** 2786-9.

```
                                         10        20        30        40        50        60
                                 ....|....| ....|....| ....|....| ....|....| ....|....| ....|....|
Arabidopsis thaliana             -MATTLQKLS SQIH--RLSP FTRSL---IV RTSATS---- --APSPSLGS KKVSDRIVKL  48
Brassica rapa                    -MATNLLKLS SQIR--RLSP LTRSL---TV RTSATS---- -------TGS KKVSDRIVKL  43
Capsella rubella                 -MATTLQKLS SQIH--RLSP FVRSL---TV RTSATS---- --APSPSLGS KKVSDRIVKL  48
Thlaspi arvense                  -MATHLQKLS SQIH--RLSP FARSV---II RTSATSAPVS -VAPSPSSGT RKVADRIVKL  53
Solanum lycopersicum             MAIAYLRRFA SRIA--RNTS LSSSFGSSIA RSSASS---- ----SSPSAS AKVADRIVKL  50
Selaginella moellendorffii       -MADLL---P VQVR------ ---------- ---------- ----QPGDSL PRVDTRIVTA  26
Physcomitrella patens            MAMAAAQSLR GSRR--HLLS YVRYGGSIAT QSESFV---- -----VRQGS AKVSDRVLEM  49
Sphagnum fallax                  -MAMAMVAGR RCLQ--RAVR HAATAAAPAV QVTQES---- --TRREAGRS AKVSDRMLEM  51
Marchantia polymorpha            -MATALRRL- VQVR------ --------FA HTAVAS---- -VSSAIPRSA PKVLESTVRL  39
Volvox carteri                   -MA--LRALG RQLRSLNLIP VVQPR--RFF GASAHHDDEH -EEEHGPAQT PTVFDKLVEI  54
Dunaliella salina                MLKAAGASLR RALAAVSTSP LESSS---RL IASSSAVQGG ADEHHGYEQT PTVFDRLVTI  57
Chlamydomonas reinhardtii        -MA--LRALG RQLRSLNLLP AVQPA--RFF GAGAHHDDEH EEEEHGPAQT PTVFDKLVEV  55
Polytomella sp.                  -MA-LLRALA KPLRSLQAVS SVAQVSLRQF GAASHHDDHH -DDHDHYTPP KTVFEDTITI  57

                                         70        80        90       100       110       120
                                 ....|....| ....|....| ....|....| ....|....| ....|....| ....|....|
Arabidopsis thaliana             SAI-DPDGYK QDIIGLSGQT LLRALTHTGL IDPASHRLDD IEACS---AE CEVQIAEEWL  104
Brassica rapa                    SAI-DADGDK KDIIGLSGQT LLRALTHTGL IDPLSHRLED IDACS---AE CEVQIAEEWL  99
Capsella rubella                 SAI-DPDGYK QDIIGLSGQT LLRALTHTGL IDPASHRLDD IEACS---AE CEVQIAEEWL  104
Thlaspi arvense                  SAI-DADGYK KEIIGLSGQT LLRALTHTGL IDPASHRLDD IEACS---AE CEVQIAEEWL  109
Solanum lycopersicum             SAV-DPDGNK RDVVGLSGQT LLKALTNQGL IDPDSHRLED IDACS---AE CEVHIAQEWL  106
Selaginella moellendorffii       IAV-DEDGRR HTVRALAGQS LFRALVNAGL RLHLSHQL-D TSCYG---YV CRVSISDEWR  81
Physcomitrella patens            TAI-DEDGQR HQIKGLTGHT LLRTLVERGL FDPERHRLEN INACG---GE CEVSIANEWL  105
Sphagnum fallax                  TAI-DEDGER HVVKGLTGHT LLRTLVQSGL FDPERHRLED ITACN---GE CEVSIANEWL  107
Marchantia polymorpha            TAIAEENGSR HAIRGLVGQT LLKTLVRSGL IEGESHRTED LTQCG---AE CEVSVAGEWL  96
Volvox carteri                   TVV-DMNGVL HKVRGLAGQT LAQALIEYGF --PATYFFPN MGFYTQHIPD AHVYVPKDYW  111
Dunaliella salina                NVV-DLDGKR RSVRAVVGQT LAQVLVEAGY --PRTYFFPN MGFYTQHIVD AHVFIPQDYW  114
Chlamydomonas reinhardtii        TVV-DMNGIR HRVRGLQGQS LAQALVEYGF --PDTYFFPN MGFYTQHIVD AHVFVPKEFW  112
Polytomella sp.                  NVL-DYDGKK HAVKALIGTP LNKALVEYGF --SSTYFFPN MGYYTQHISD AHVFIPEEYW  114
                                                                                *          *
                                        130       140       150       160       170       180
                                 ....|....| ....|....| ....|....| ....|....| ....|....| ....|....|
Arabidopsis thaliana             EKLP--PRTY DEEYVL-KRS SRSRILN--- KHSRLGCQVV LTQELQGMVV AVPEAKPWDI  158
Brassica rapa                    EKLP--PRTY DEEYVL-KRN SRSRVLN--- KHSRLGCQVV LTQELQGMVV AVPEAKPWDI  153
Capsella rubella                 EKLP--PRSY DEEYVL-KRS SRSRVLN--- KHSRLGCQVV LTQELQGMVV AVPEAKPWDI  158
Thlaspi arvense                  EKLP--TRTY DEEYVL-RRN SRSRVLN--- KHSRLGCQVV LTQELQGMVV AVPEAKPWDI  163
Solanum lycopersicum             EKLP--PASY DEKYVL-KRN SRARVLN--- KHSRLGCQVV LSQELQGMVV ALPEPKPWDI  160
Selaginella moellendorffii       ARIP--EPTE DELHVL-KHH FTPPDFD--- PSIRCSCKIK LSDKINGIVV ALVPEKCWDF  135
Physcomitrella patens            DKLP--PRSE DELEVL-KDK THAKKAD--- PHARLGCQIV LEPELEGMVV SIAEEKPWRT  159
Sphagnum fallax                  VKLP--PRTE DELDVL-QAK TFSKLAD--- PHSRLGCQII LEPELEGMVV SIAEEKPWRT  161
Marchantia polymorpha            EKLP--PRSE DEMEILEKHQ PKGQAVD--- THTRLACQIV LDRKLDGMAF AIPEARPWMS  151
Volvox carteri                   GKMPNIDPES EEGGAV-KRM FRDIVQDYQR DTSYFASYIT LGPELNGMTV GIGPIKPWIL  170
Dunaliella salina                SKIRNYKEDS DEADAI-KRT FRDMIQDYAK STSYLASYID IVPEFNNMTI GIGPVKPWVL  173
Chlamydomonas reinhardtii        GKVQNVDPES DDGLAV-KRM FRDIVQDYQR DTSFFASYIT LGAEHNGMTV GIGPIKPWIL  171
Polytomella sp.                  KYVENVDLKT DDAEAI-KLM FKLVVQDYQR ETSFFASYLT LNKEMDNMTI GFGPIKPWHI  173
                                                                      *
                                        190       200       210
                                 ....|....| ....|....| ....|....|
Arabidopsis thaliana             P--------- ---------- ----------  159
Brassica rapa                    P--------- ---------- ----------  154
Capsella rubella                 P--------- ---------- ----------  159
Thlaspi arvense                  P--------- ---------- ----------  164
Solanum lycopersicum             P--------- ---------- ----------  161
Selaginella moellendorffii       T--------- ---------- ----------  136
Physcomitrella patens            L--------- ---------- ----------  160
Sphagnum fallax                  L--------- ---------- ----------  162
Marchantia polymorpha            N--------- ---------- ----------  152
Volvox carteri                   HSDWAFAGVH DSKAKQFDK- ---PTVEIWG  196
Dunaliella salina                HSEWAFNGVH DDPTPQFKE- ---PTTEIWG  199
Chlamydomonas reinhardtii        HSEWAFAGVH DSRNKQFDK- ---PTVEIWG  197
Polytomella sp.                  TPKWSFNGHH NVKDRMFDRL ETGPFIE---  200
```

**Supplemental Figure S2. Amino acid sequence alignment of selected plant and algal mFDX-like proteins.** The three remaining cysteines known to serve as [2Fe-2S] cluster ligands in regular FDXs are pointed by asterisks. Shaded areas indicate at least 80% amino acid identity (black) or similarity (grey). The UniProt accession numbers are: *Arabidopsis thaliana*: Q9SRR8, *Brassica rapa*: M4ELI3, *Capsella rubella*: R0I0R4, *Solanum lycopersicum*: A0A3Q7GPB6, *Selaginella moellendorffii*: D8RBK4, *Physcomitrella patens*: A0A2K1L3S2, *Marchantia polymorpha*: A0A2R6WIY0, *Volvox carteri*: D8TW15, *Chlamydomonas reinhardtii*: Q6QAY1. The sequences absent from UniProt were retrieved from the following sources: Phytozome: *Thlaspi arvense*: Thlar.0128s0036, *Sphagnum fallax*: Sphfalx07G032200, *Dunaliella salina*: Dusal.0767s00002 and NCBI: *Polytomella sp.*: 7AR9_O.

*

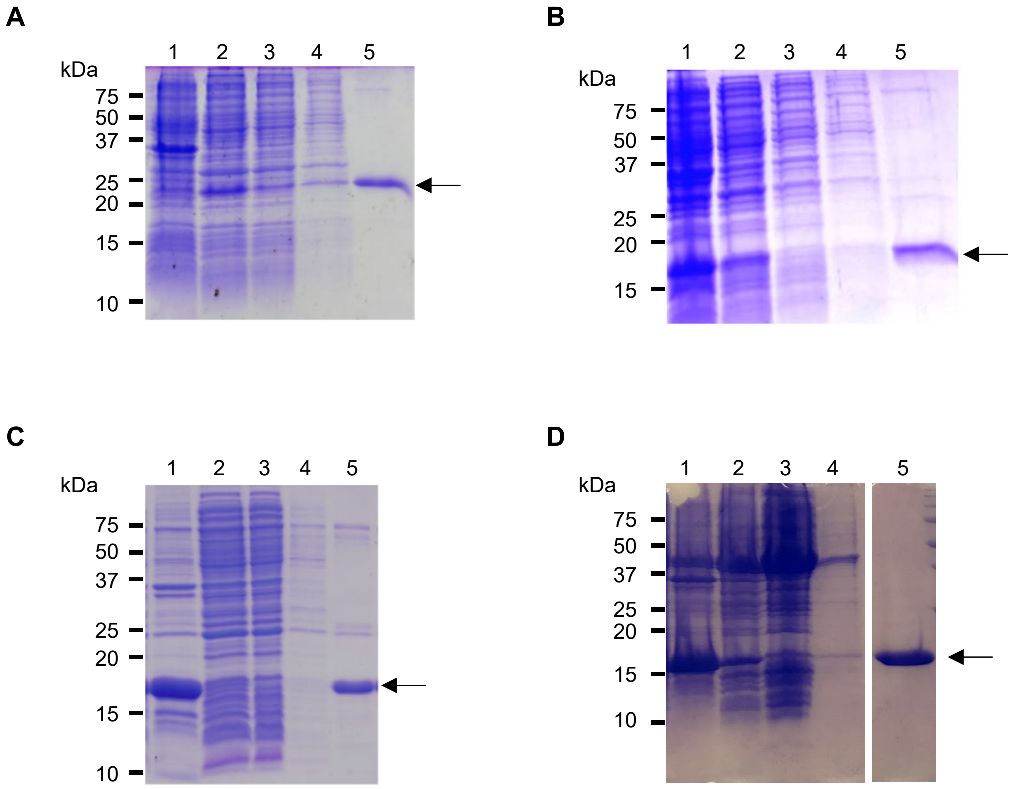


**Supplemental figure 3. Purification steps and purity degree of recombinant proteins.**

Fractions obtained during aerobic protein purification of tagged proteins were analyzed on 15% SDS-PAGE. 1. insoluble fraction, 2. soluble fraction, 3. flowthrough fraction, 4. washing fraction and 5. elution fraction. A. mFDX1, B. mFDX2, C. mFDX-like, D. mFDX-like L85C. The arrows indicate the bands corresponding to the expected size of each proteins. In D, the gel was cut because of an intermediate lane but it is the same gel.

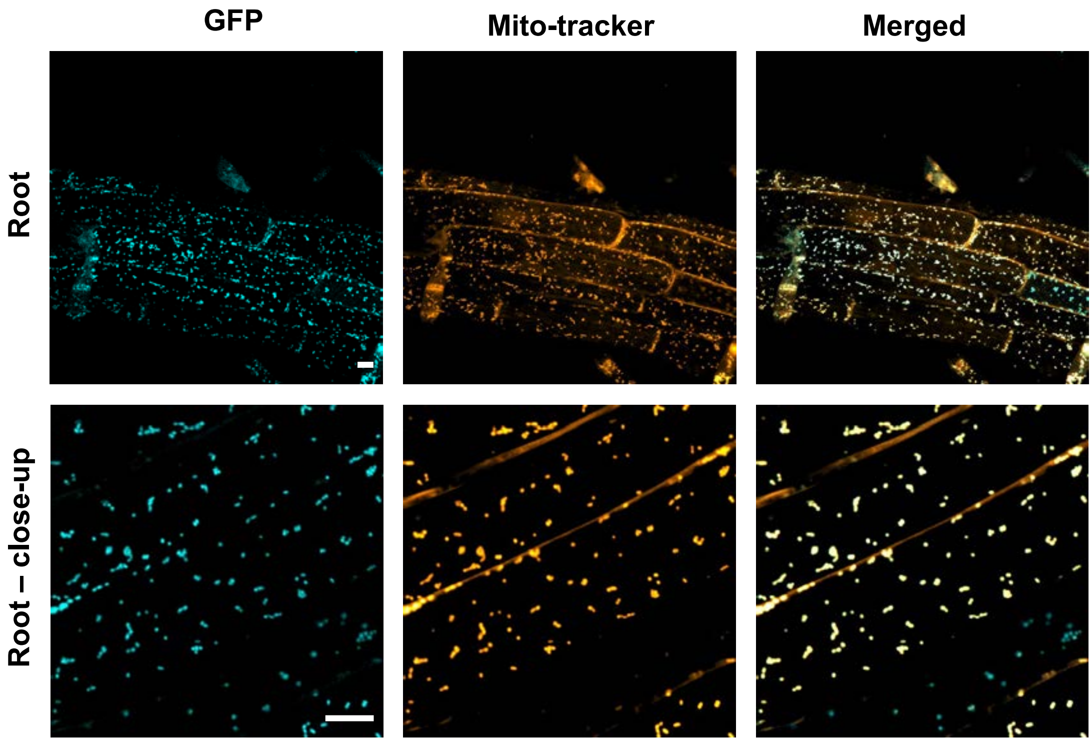


**Supplemental Figure S4. Subcellular localization of AtmFDX-like in Arabidopsis.** Expression of *promoter-mFDX-like::mFDX-like::GFP* in root cells from a 14-day-old seedling. GFP, cyan; MitoTracker, orange. Scale bar: 10 µm.

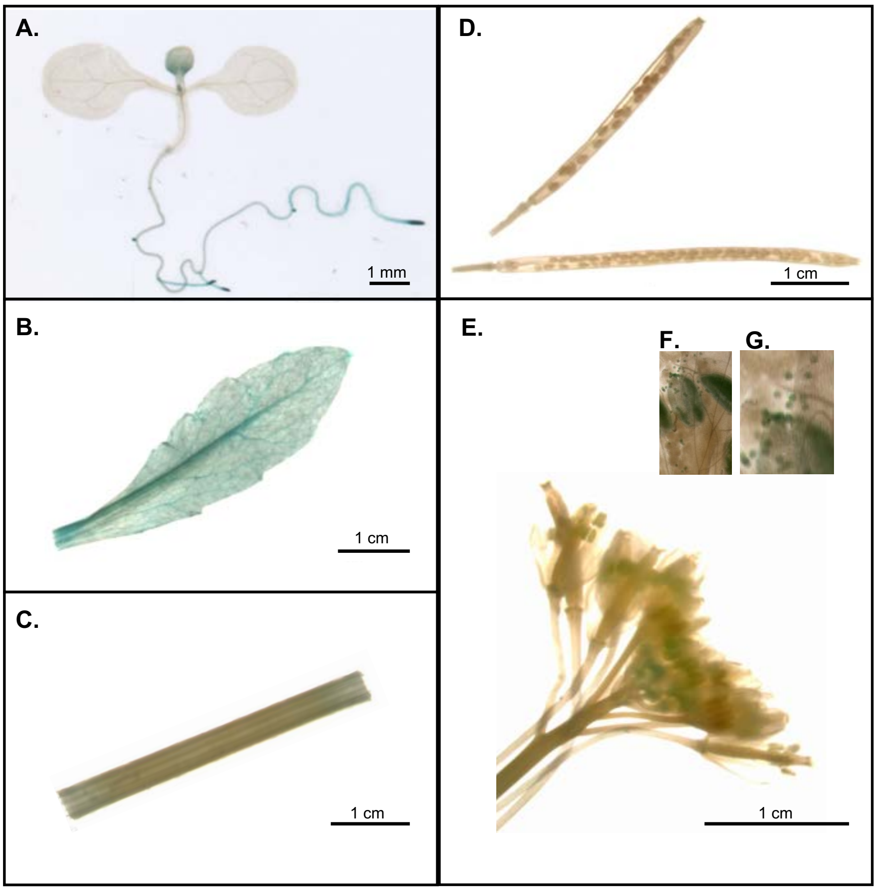


**Supplemental Figure S5. Histochemical localization of GUS in transgenic *Arabidopsis thaliana* plants containing the *promoter-mFDX-like::mFDX-like::GUS.*** GUS activity was assayed in **(A)** 7-day-old seedling, 6-week-old **(B)** leaf, **(C)** stem, **(D)** siliques, **(E)** flowers, **(F)** anthers and **(G)** pollen grains.

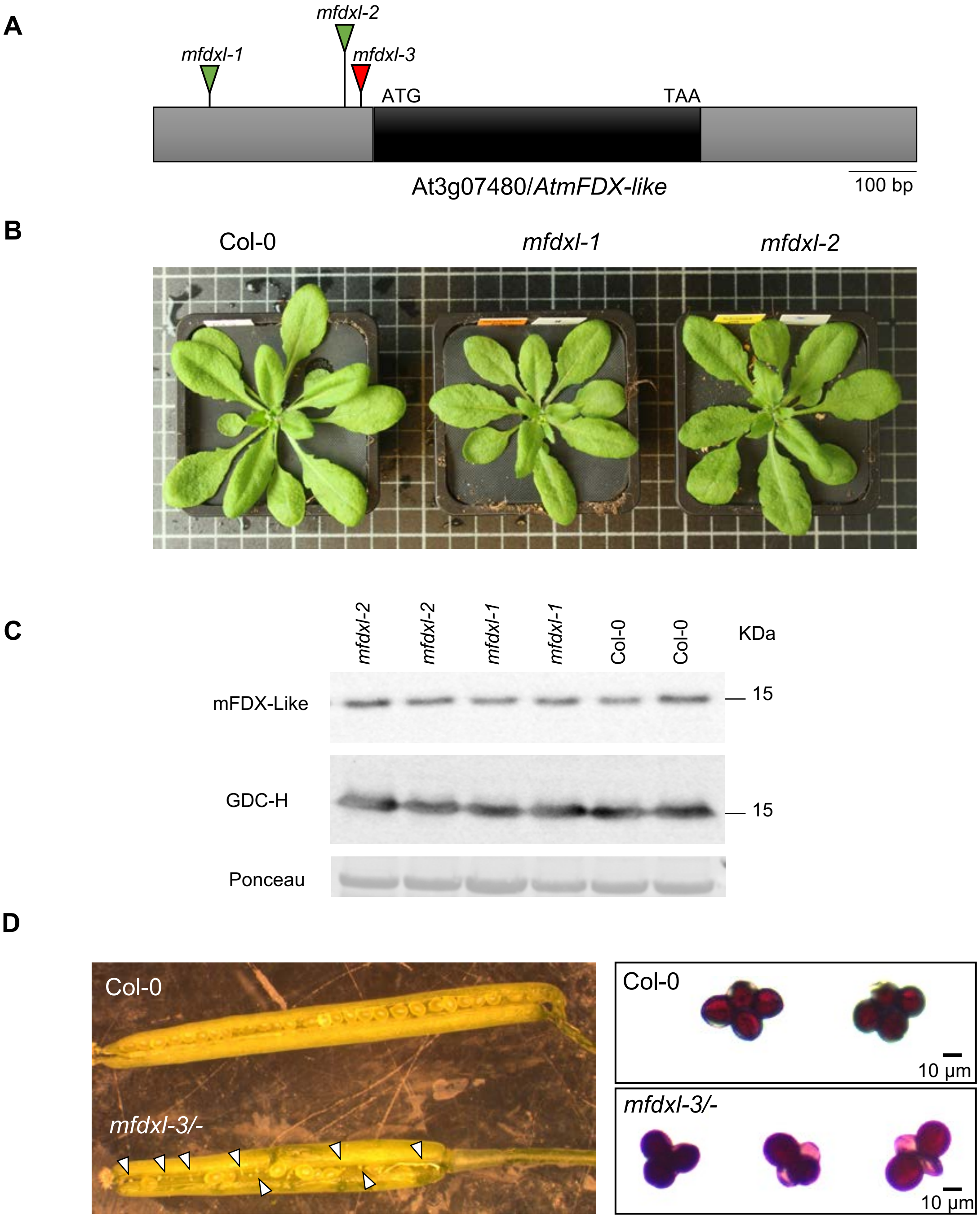


**Supplemental Figure S6. Genetic analysis and characterization *mfdx-like T-DNA insertion* mutants from *Arabidopsis thaliana*.**

(A) Gene structure of the *mFDX-like* gene, At3g07480. Exons are represented in black, both 5' and 3' untranslated (UTRs) regions are shown as grey rectangle. Positions of T-DNA insertions of the mutant lines are represented by triangles (green triangles indicate that homozygous mutants were retrieved whereas the red triangle indicates that no homozygous mutants were obtained). *mfdxl-1*: SALK_015333; *mfdxl-2*: WiscDsLoxHs204_01A; *mfdxl-3*: SAIL_37_H11. (B) Growth phenotypes of 24 day-old wild-type Col-0 (WT), *mfdxl-1* and *mfdxl-2* plants grown under long-day conditions. (C) Abundance of mFDX-like protein levels determined by western-blot. These representative western-blots were performed from mitochondria-enriched extracts isolated from leaves. 20 µg of proteins was loaded per lines. Levels of H-proteins and Ponceau S stain was used to confirm equal loading and transfer. (D) Siliques from a wild-type (top) and mfdxl-3/- (bottom) were opened and immature seeds are shown. Mature pollen grains were collected from wild-type and heterozygous *mfdxl-3* plants and viability was tested by using Alexander's stain. Cytoplasm of viable pollen grains was colored in purple whereas dead cells appeared transparent and shriveled.

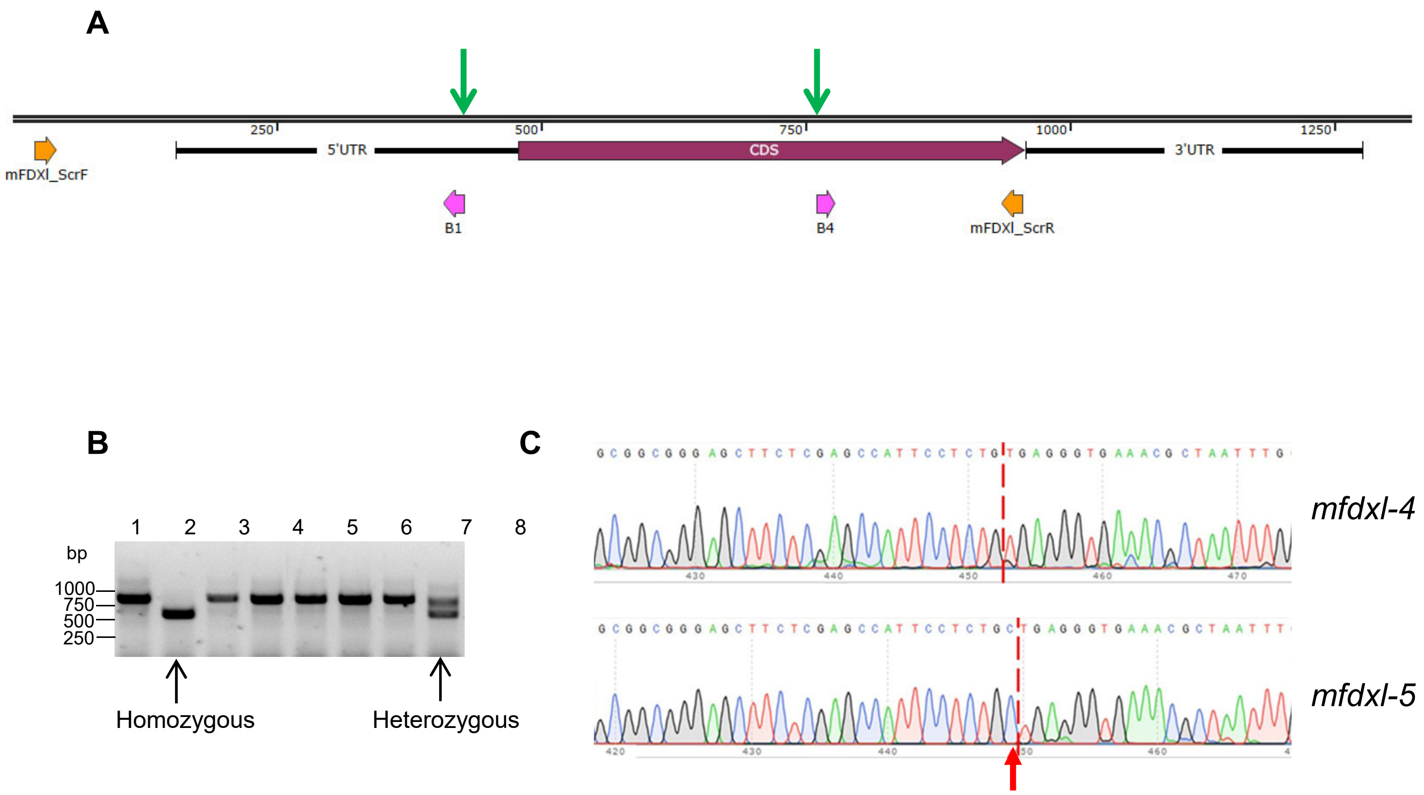


**Supplemental Figure 7. CRISPR/Cas9 strategy to obtain deletion mutants.**

A. Schematic representation of the gene *At3g07480* coding for mFDX-like. The protospacer sequences of the two guide RNAs (B1 and B4) used to create the deletions are indicated with pink arrows. The expected Cas9 cut sites are indicated with green arrows. The sequence between the two cuts is lost after repair of the DNA. The position of the primers used to screen the mutant lines is indicated with orange arrows. Numbers indicate the number of nucleotides from the left end of the scheme. B. Eight T2 plants were screened by PCR using the primers mFDXl_ScrF and mFDXl-ScrR. The WT PCR product has a size of 934 bp, the deletion product has an expected size of ca. 600 bp. No signal is detected at 934 bp in plant 2, this plant is thus considered homozygous. C. Sequence chromatograms obtained by sequencing the 600 bp PCR product of two homozygous lines using the primer mFDXl_ScrR. The mutant *mfdxl-4* shows the expected repair site (indicated with a dashed red line) whereas the mutant *mfdxl-5* contains an additional C (indicated with a red arrow) at the repair site (dashed red line).

**Supplemental Table S1**: **UniProt accession numbers of the proteins used for the phylogenetic study.**

Eukaryotic groups are based on (Keeling and Burki 2019). For each organism, the different FDXs were identified using PSI-BLAST using the Arabidopsis mFDXs as baits. The coresponing UniProt accession is given. The chloroplastic FDXs are indicated in green

<table>
<tr><th>Group</th><th>Class</th><th>Organism</th><th>Name</th><th>Uniprot</th></tr>
<tr><td rowspan="2">Amoebozoa</td><td>Discosea</td><td>Acanthamoeba castellanii</td><td>AcFDX</td><td>L8GR90</td></tr>
<tr><td>Dictyostelia</td><td>Dictyostelium discoideum</td><td>DdFDX</td><td>Q55GW1</td></tr>
<tr><td></td><td>Apusomonadea</td><td>Thecamonas trahens</td><td>ThFDX</td><td>A0A0L0DMW7</td></tr>
<tr><td rowspan="3">Opisthokonta</td><td rowspan="2">Chordata</td><td rowspan="2">Homo sapiens</td><td>HsFDX1</td><td>P10109</td></tr>
<tr><td>HsFDX2</td><td>Q6P4F2</td></tr>
<tr><td>Ascomycota</td><td>Yarrowia lipolytica</td><td>YlFDX</td><td>Q6CFZ5</td></tr>
<tr><td rowspan="15">Archaeplastida</td><td rowspan="9">Tracheophyta</td><td rowspan="9">Arabidopsis thaliana</td><td>AtcFDX1</td><td>O04090</td></tr>
<tr><td>AtcFDX2</td><td>P16972</td></tr>
<tr><td>AtcFDX3</td><td>Q9ZQG8</td></tr>
<tr><td>AtcFDX4</td><td>Q9FIA7</td></tr>
<tr><td>AtcFDXC1</td><td>O23344</td></tr>
<tr><td>AtcFDXC2</td><td>Q9C7Y4</td></tr>
<tr><td>AtmFDX1</td><td>Q9M0V0</td></tr>
<tr><td>AtmFDX2</td><td>Q8S904</td></tr>
<tr><td>AtmFDXlike</td><td>Q9SRR8</td></tr>
<tr><td rowspan="3">Chlorophyta</td><td rowspan="3">Chlamydomonas rheinhardtii</td><td>CrcFDX</td><td>Q2HZ24</td></tr>
<tr><td>CrFDX1</td><td>A8JHR1</td></tr>
<tr><td>CrNUOP3</td><td>Q6QAY1</td></tr>
<tr><td rowspan="3">Rhodophyta</td><td rowspan="3">Cyanidioschyzon merolae</td><td>CmcFDX1</td><td>M1V5H0</td></tr>
<tr><td>CmcFDX2</td><td>Q85FT5</td></tr>
<tr><td>CmcFDX2</td><td>Q85FT5</td></tr>
<tr><td rowspan="3">Cryptista</td><td rowspan="3">Cryptophyta</td><td rowspan="3">Guillardia theta</td><td>GtcFDX</td><td>O78510</td></tr>
<tr><td>GtFDX1</td><td>L1IPS0</td></tr>
<tr><td>GtFDX2</td><td>L1J5A6</td></tr>
<tr><td></td><td>Picozoa</td><td>Picomonas judeaskeda</td><td colspan="2">no ferredoxins identified</td></tr>
<tr><td></td><td>Malawimonadea</td><td>Malawimonas jakobiformis</td><td colspan="2">no ferredoxins identified</td></tr>
<tr><td rowspan="3">Haptista</td><td rowspan="3">Haptophyta</td><td rowspan="3">Emiliania huxleyi</td><td>EhcFDX</td><td>R1DCW6</td></tr>
<tr><td>EhFDX1</td><td>R1DSP5</td></tr>
<tr><td>EhFDX2</td><td>R1D133</td></tr>
<tr><td rowspan="11">Stramenopila</td><td rowspan="5">Eustigmatophycea</td><td rowspan="5">Nannochloropsis gaditana</td><td>NgcFDX1</td><td>W7U5Y2</td></tr>
<tr><td>NgcFDX2</td><td>W7TUL4</td></tr>
<tr><td>NgFDX1</td><td>W7TQA0</td></tr>
<tr><td>NgFDX2</td><td>K8YUL7</td></tr>
<tr><td>NgFDX3</td><td>W7TRL6</td></tr>
<tr><td rowspan="4">Diatoms</td><td rowspan="4">Thalassiosira oceanica</td><td>TocFDX</td><td>K0RP06</td></tr>
<tr><td>ToFDX1</td><td>K0TE65</td></tr>
<tr><td>ToFDX2</td><td>K0SI29</td></tr>
<tr><td>ToFDX3</td><td>K0RID3</td></tr>
<tr><td rowspan="2">Oomyceta</td><td rowspan="2">Phytophtora infestans</td><td>PiPDX1</td><td>D0MSF6</td></tr>
<tr><td>PiPFD2</td><td>A0A833RND4</td></tr>
</table>

| Group | Class | Organism | Name | Uniprot |
|---|---|---|---|---|
| Alveolata | Apicomplexa | *Toxoplasma gondii* | TgFDX1 | B6KFF5 |
| | | | TgFDX2 | B9Q746 |
| | Ciliophora | *Tetrahymena thermophila* | TtFDX1 | Q22VV0 |
| | | | TtFDX2 | I7M2B0 |
| | | | TtNDUFX | Q22W11 |
| Rhizaria | Foraminifera | *Reticulomyxa filosa* | RfFDX1 | X6PDJ4 |
| | | | RfFDX2 | X6NJJ3 |
| Discoba | Kinetoplastida | *Trypanosoma brucei* | TbFDX1 | Q57VT5 |
| | | | TbFDX2 | Q584K7 |
| | Euglenoidea | *Euglena viridis* | EvcFDX | P22341 |
| Metamonada | Diplomonadida | *Giardia intestinalis* | GiFDX | E2RU02 |
| CRuMS | Mantomonada | *Mantomonas plastica* | no ferredoxins identified | |

KEELING, P. J. & BURKI, F. 2019. Progress towards the Tree of Eukaryotes. *Curr Biol,* 29**,** R808-R817.

**Supplemental Table S2.** Primers used in this study

| Primer name | Primer sequence | Use |
|---|---|---|
| AtmFDX1for | CCCCCCCCCATATGTCCTCTGAGAATGGTGAT | Cloning in pET vectors |
| AtmFDX2for | CCCCCCCCCATATGACCTCTGAGAAAGGTGGC | Cloning in pET vectors |
| AtmFDX1/2rev | CCCCGGATCCCTAATGAGGTTTTGGAACAAACCC | Cloning in pET vectors |
| AtmFDX-like for | CCCCCCCCATATGGGATCGAAGAAAGTCTCT | Cloning in pET vectors |
| AtmFDX-like L85C for | CCAGCATCGCATAGAT**GT**GATGACATCGAGGCT | Cloning in pET vectors |
| AtmFDX-like rev | CCCCGGATCCTTACGGAATATCCCAAGG | Cloning in pET vectors |
| AtmFDX-like L85C rev | AGCCTCGATGTCATC**AC**ATCTATGCGATGCTGG | Cloning in pET vectors |
| SALK015333RP | ATGTCCTGCTTGTATCCATCG | Genotyping SALK_015333 |
| SALK015333LP | ATAGTCCCCGTAATATTGGCG | Genotyping SALK_015333 |
| SALK_LBb1.3 | ATTTTGCCGATTTCGGAAC | Genotyping SALK_015333 |
| SAIL434RP | ACACAACTTGGCAACCAAGAC | Genotyping SAIL_434_E06 |
| SAIL434LP | AAACGCGATTTGAACAATACG | Genotyping SAIL_434_E06 |
| SAIL_LB3 | TAGCATCTGAATTTCATAACCAATCTCGATACAC | Genotyping SAIL_434_E06 |
| Wisc204_RP | AAGGCTTAGCTTCAGGGACAG | Genotyping WiscDsLoxHs204_01A |
| Wisc204_LP | AAACGCGATTTGAACAATACG | Genotyping WiscDsLoxHs204_01A |
| Wisc_LB | TGATCCATGTAGATTTCCCGGACATGAAG | Genotyping WiscDsLoxHs204_01A |
| oJF212 | tgccAAGCTTcgacttgcct | Building of pJF1033 |
| oJF215 | attcACTAGTataaccatggtattgg | Building of pJF1033 |
| oJF398 | accaGGTCTCaATTGAAACCGGAAAAGTGAGTGAgtt ttagagctagaaatagcaag | Building of pJF1120 |
| oJF399 | tggtGGTCTCtAAACCTGCGATCTGAACCTCGCACaa tctcttagtcgactctacc | Building of pJF1120 |
| mFDXI_ScrF | AACAGACCAATGAACGATAGCC | Screening of CRISPR lines |
| mFDXI_ScrR | CGGAATATCCCAAGGCTTAG | Screening of CRISPR lines |
| Cas9_F | GTGCCAGCTGCTGACAAGAAG | Detection of Cas9 in the deletion lines |
| Cas9_R | AGAGCCTAGCGGACAGGATAGC | Detection of Cas9 in the deletion lines |
| mFDXI_GG_for | GGGGACAAGTTTGTACAAAAAAGCAGGCTTCCCA CAAGAATAGCTCTTGCGC | Cloning in pGWB3 and pGWB4 |

| | | |
|---|---|---|
| mFDXI_GG_rev | GGGGACCACTTTGTACAAGAAAGCTGGGTCCGGAATATCCCAAGGCTT | Cloning in pGWB3 and pGWB4 |
| mFDXI_Com_F | GGGGACAAGTTTGTACAAAAAAGCAGGCTTCATGGCGACGACTCTTCAG | Cloning in pGWB2 |
| mFDXI_Com_R | GGGGACCACTTTGTACAAGAAAGCTGGGTCTTACGGAATATCCCAAGGCTT | Cloning in pGWB2 |
